# Engineered prime editors with minimal genomic errors

**DOI:** 10.1101/2024.08.02.606370

**Authors:** Vikash P. Chauhan, Phillip A. Sharp, Robert Langer

## Abstract

Prime editors make programmed genome modifications by writing new sequences into extensions of nicked DNA 3’ ends. These edited 3’ new strands must displace competing 5’ strands to install edits, yet a bias toward retaining the competing 5’ strands hinders efficiency and can cause indel errors. Using rational design of the constituent Cas9-nickase to reposition prime editor nicks, we discovered that competing 5’ strands are destabilized to favor the edited 3’ new strands. We exploit this mechanism to engineer efficient prime editors with strikingly low indel errors. Combining this error-suppressing strategy with the latest efficiency-boosting architecture, we design a next-generation prime editor (vPE). Compared with previous editors, vPE features comparable efficiency yet up to 60-fold lower indel errors, enabling edit:indel ratios as high as 465:1.

**One Sentence Summary:** Prime editors designed with repositioned DNA breaks nearly eliminate undesired genome editing errors

## Main Text

Prime editors are advanced CRISPR tools that enable replacement of targeted DNA with programmed sequences (*1*). A prime editor is comprised of a Cas9 nickase (Cas9n) fused to a reverse transcriptase (RT) and paired with an extended guide RNA (pegRNA) that encodes both the genomic target sequence and intended edit (*2*). Editing initiates with the prime editor binding its genomic target and forming a single-strand DNA break (nick) (Fig. 1A). The nicked 3’ DNA end is released to anneal to the pegRNA template, priming the RT to write the template sequence into an extension of the 3’ DNA end. This resulting edited 3’ new strand can displace the competing 5’ strand to install the intended edit. This process can be adapted to a wide variety of edit types, including substitutions, insertions, and deletions (*2–8*). Significant efforts have been made to increase prime editing efficiency, including inhibition of cellular mismatch repair (MMR) (*9*, *10*), stabilization of pegRNAs (*11–13*), and engineering the RT domain (*9*, *14–16*). These advances have led to highly effective and versatile prime editing systems.

**Figure 1.**
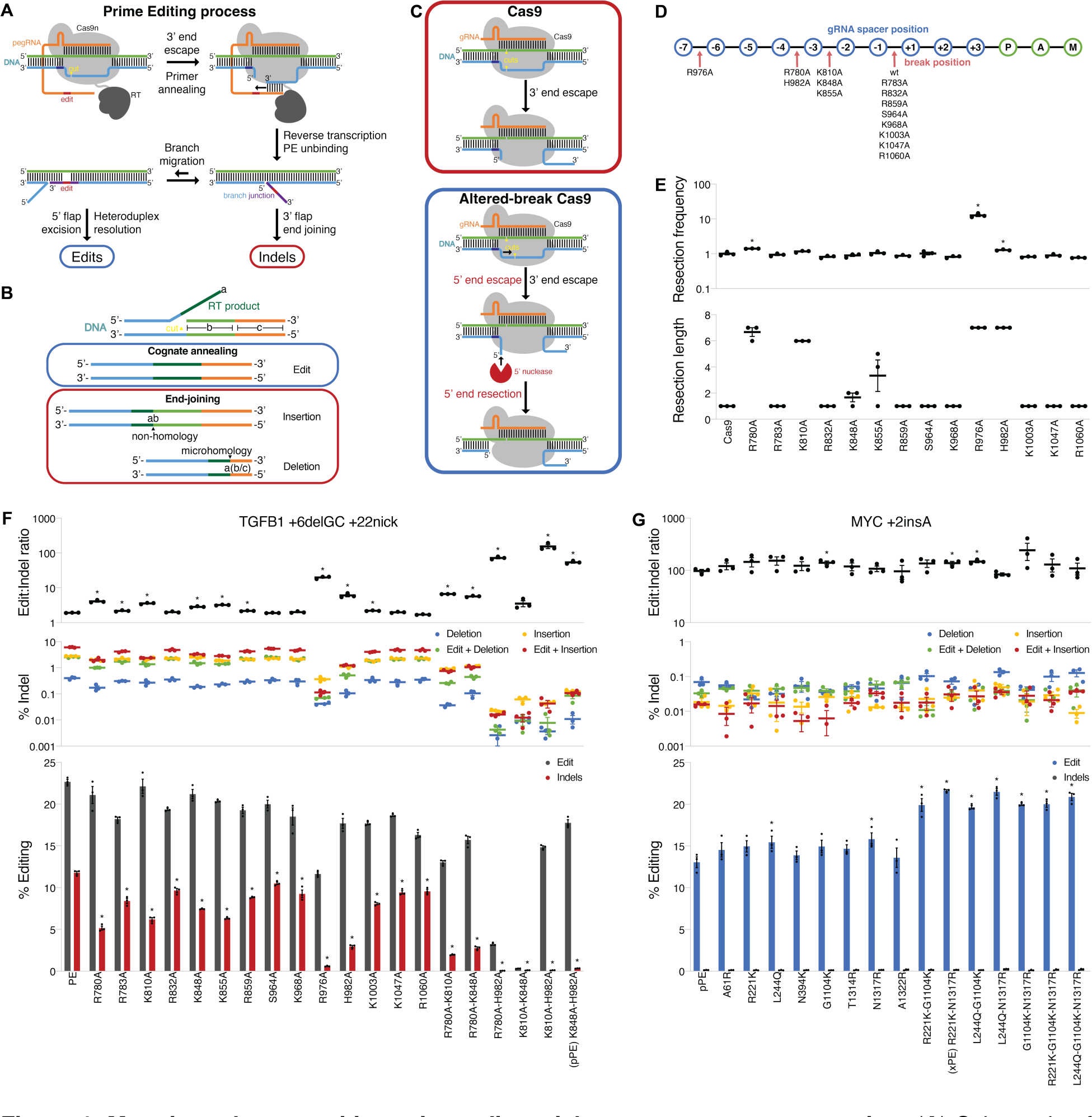
Mutations that reposition prime editor nicks suppress error generation. (**A**) Schematic of the prime editing process, depicting two competing structures that can produce edits or indel errors. (**B**) Schematic of mechanisms through which indel errors are produced by end-joining of edited 3’ new strands. (**C**) Model depicting how Cas9 variants with different break structures are proposed to either protect nontarget strand 5’ ends (top) or promote their resection (bottom). (**D**) Positions of most frequent shifted nontarget strand nicks, from analysis of public data (*21*). (**E**) The degree (bottom) and frequency (top) of nontarget strand 5’ end resection for different Cas9 variants, from analysis of public data (*21*). (**F**) Screen of engineered PE variants to suppress indel errors with quantification of edit and indel frequencies (bottom), indel classes (middle), and edit:indel ratios (top). (**G**) Screen of engineered pPE variants to enhance efficiency with quantification of edit and indel frequencies (bottom), indel classes (middle), and edit:indel ratios (top). * indicates p < 0.05 for comparisons to Cas9 in **E**, to PE in **F**, and to pPE in **G**. All data were analyzed by deep sequencing and represent means of *n* = 3 independent replicates with standard errors.

A key remaining challenge is the elimination of errors that occur as byproducts of prime editing. These errors are insertion-deletion (indel) mutations generated in lieu of the intended edit within a fraction of targeted cells, resulting in DNA sequences that are unpredictable and possibly deleterious (*9*, *17*). Previous work has identified major drivers of indel error formation, though the mechanisms have not been fully elucidated. First, the prime editor may extend the edited 3’ new strand past the pegRNA template and into the scaffold (*2*, *9*). This can be addressed by recoding the pegRNA scaffold to limit its homology with the genomic sequence (*9*). Second, errant double-strand breaks (DSB) can be generated, sometimes as a consequence of MMR converting nicks into DSBs, and can induce indels consistent with DSB repair (*9*, *18*). These indels can be addressed through inhibition or avoidance of MMR (*9*, *10*). Third, the edited 3’ new strand generated by the prime editor can end join at unintended positions (*9*). This often produces large deletions or tandem duplication-like insertions, and there are presently no strategies for addressing these errors.

In the prime editing process, the edited 3’ new strand is disfavored in displacing the competing 5’ strand against annealing of the edited 3’ new strand can limit editing efficiency and promote errors (Fig. 1B). It is known that the 3’ end of a nicked DNA substrate can escape from bound Cas9 complex whereas the shorter 5’ end remains stably bound (*19*, *20*). We hypothesized that repositioning the nick to lengthen the 5’ end might release it from Cas9 and enable its resection by cellular nucleases (Fig. 1C). We recently discovered that Cas9 can be engineered to alter nick positioning (*21*), which encouraged our exploration here of whether competing 5’ strands formed by prime editors can be destabilized by lengthening them. In this study, we examined mutations that reposition Cas9 nicks to possibly enable 5’ end resection. Using these mutations, we engineered prime editors to discover an unexpectedly large influence of nick positioning on indel error generation. Through rational design, we created efficient prime editors that rarely produce indel errors.

### Altering prime editor nick positioning suppresses indel error generation

To determine whether the 5’ ends at Cas9-induced nicks can be destabilized, we characterized 5’ end resection for engineered Cas9 variants. We recently screened *S. pyogenes* Cas9 Alanine substitutions in the DNA binding clefts and identified mutations that reposition the nontarget strand nick (Fig. 1D) (*21*). Therein we generated a sequencing dataset where nearby DSB pairs were created using wild-type Cas9 and fourteen mutants, and here we analyzed the dataset to quantify 5’ end resection. In this assay paired DSBs produce junctions between the two cuts, and we reasoned that additional deletions would indicate the degree and frequency of nontarget strand 5’ end resection (fig S1A). We quantitated the median length and frequency of such deletions as metrics of resection for these fifteen Cas9 variants (Fig. 1E and fig. S1B). Our analysis indicated that wild-type Cas9 produced minimal resection, consistent with nicked 5’ end protection. Junction sequences revealed that wild-type Cas9 mostly produced 1bp deletions but rarely larger deletions, suggesting minimal resection (fig. S1C). We similarly observed minimal resection for several Cas9 variants without altered nick positions. In contrast, mutations that repositioned the nick by 2nt (K810A, K848A, and K855A) led to increased resection. Further, mutations that shifted the nick position 3nt Junction sequences for R976A and H982A revealed dominant deletions several bp in length indicating significant resection (fig. S1C). These findings supported a model where repositioning the nontarget strand nick to lengthen the 5’ end likely rendered it accessible to nuclease degradation.

Since repositioning the nontarget strand nick promoted 5’ end resection, we explored whether prime editors with nick-repositioning Cas9n mutations might show enhanced editing. We began with PEmax, composed of Cas9n(H840A) with R221K-N394K mutations (*9*). For direct comparison to the screened Cas9 variants, we reverted these to R221-N394 and named this prime editor PE. We evaluated PE using small edits at six loci (*EMX1*, *GFP*, *KRAS*, *MYC*, *STAT1*, and *TGFB1*) in HEK293T cells and observed modestly reduced efficiency versus PEmax (fig. S2A). To probe how nick-repositioning mutations affect editing outcomes, we introduced the fourteen mutations from our Cas9 screen into PE. We screened these PE variants targeting edits at the *TGFB1* and *KRAS* loci in HEK293T cells with the pegRNA+nicking gRNA (ngRNA) mode (Fig. 1F and fig. S2B). Several PE variants (R780A, K810A, K848A, R976A, and H982A) decreased indel errors up to 20-fold, with improved edit:indel ratios up to 10-fold. We observed similar suppression of four different indel classes (deletions and insertions, without or with the edit). Notably, there was a strong correlation between nick repositioning, 5’ end resection, and indel error reductions for different mutations. This supported a connection between competing 5’ strand resection and indel error suppression, suggesting this as a new mechanism for enhancing prime editing fidelity.

Considering indel errors were greatly suppressed by mutations that reposition nicks, we combined these mutations to create double-mutant PE variants and similarly tested them (Fig. 1F and fig. S2B). These variants demonstrated greatly reduced indel errors, up to 118-fold lower than PE. One variant, K848A- H982A, nearly eliminated errors, reducing them 36-fold versus PE and improving the edit:indel ratio 28-fold. We named this variant precise prime editor, or pPE. Comparing pPE to PEmax across six loci (*CXCR4*, *EMX1*, *GFP*, *MYC*, *STAT1*, and *TGFB1*) in HEK293T cells revealed consistent indel error suppression. For pegRNA-only editing versus PEmax, pPE reduced indels 7.6-fold (range 1.1 – 13-fold) and increased the edit:indel ratio 6.3-fold (range 0.4 – 10-fold) (fig. S3, A-D). These improvements versus PEmax were more dramatic for pegRNA+ngRNA editing, where pPE decreased indels 26-fold (range 7.7 – 36-fold) and improved the edit:indel ratio 20-fold (range 6.6 – 39-fold) (fig. S3, E-H). Examination of different indel classes demonstrated significant reductions of each (fig. S3, C and G). For pegRNA-only editing these reductions were 2.5- to 25-fold, and for pegRNA+ngRNA editing they were 7.8- to 28-fold. We observed similar improvements in the presence of MMR inhibition (fig. S3, I-P). Remarkably, these gains for pPE enabled edit:indel ratios of up to 361:1 (fig. S3, D and H). Therefore, the effects of altering nick positioning on error generation extended to diverse edit types and prime editing modes.

### Further design boosts efficiency and minimizes errors

While pPE suppressed errors versus PEmax, we observed a moderate reduction in editing efficiency (fig. S3). Similar reductions in Cas9 activity were associated with efforts to improve on-target specificity, expand targeting space, and alter repair outcomes (*21–26*). Accordingly, we reasoned that decreased efficiency might be addressed by incorporating mutations previously found to enhance Cas9 activity into pPE (*21*, *25–27*). We tested eight mutations that introduce charged residues near the nuclease positions in Cas9 to possibly rescue reduced activity. We screened these pPE variants using edits at the *MYC* and *STAT1* loci in HEK293T cells in the pegRNA-only mode (Fig. 1G and fig. S4A). Several mutants demonstrated increased efficiency up to 1.2-fold with modestly improved edit:indel ratios. Notably, two mutations (R221K and L244Q) are known to non-specifically increase activity for Cas9 (*27*), whereas the other two (G1104K and N1317R) were near A982 and increase Cas9 activity in the context of nearby mutations (*21*, *26*). We then assessed combinations of these mutations to identify variants with maximized efficiency (Fig. 1G and fig. S4A). We observed increased efficiency up to 1.7-fold, again with improved edit:indel ratios. Additional inclusion of N863A to reduce potential DSB formation (*18*) did not improve edit:indel ratios, suggesting that our variants caused minimal DSBs (fig. S4B). The most efficient variant, R221K-K848A-H982A-N1317R, also improved edit:indel ratios from 276:1 for pPE to 354:1. We named this variant extra-precise prime editor, or xPE.

We investigated the capabilities of these engineered prime editors to understand the mechanisms by which they improved editing. We compared xPE to PEmax across six loci (*CXCR4*, *EMX1*, *GFP*, *MYC*, *STAT1*, and *TGFB1*) in HEK293T cells. For pegRNA-only editing, we observed a decrease in indel rates from 0.59% (range 0.095 – 1.2%) for PEmax to 0.11% (range 0.072 – 0.14%) for xPE with slightly lower rates of intended edits (fig. S5, A and B). Subclassifying indels showed similar reductions for deletions without or with the intended edit, while the decrease in insertions was less for xPE versus PEmax (fig. S5C). This corresponded to broad improvements in edit:indel ratios (fig. S5D). Summarizing our findings for pegRNA- only editing, xPE decreased indels 5.0-fold (range 1.3 – 9.2-fold) and increased the edit:indel ratio 4.2-fold (range 0.7 – 8.2-fold) versus PEmax. These effects of xPE were more dramatic for pegRNA+ngRNA editing, where we observed a reduction in indels from 16% (range 14 – 19%) for PEmax to 1.8% (range 0.52 – 3.1%) for xPE (fig. S5, E and F). Analyzing indel subclasses showed that xPE decreased all indel types compared to PEmax, most dramatically for deletions (fig. S5G). These effects for xPE led to large increases in the edit:indel ratio versus PEmax (fig. S5H). For pegRNA+ngRNA editing, xPE significantly reduced indels 12.7-fold (range 5.0 – 27.5-fold) and improved the edit:indel ratio 9.4-fold (range 4.0 – 14.1-fold) versus PEmax. These improvements for xPE led to edit:indel ratios of up to 199:1 (fig. S5, D and H).

### Stabilization of pegRNAs synergizes with error-suppressing mechanisms

While xPE gained efficiency over pPE while maintaining low error rates, we still observed slightly lower efficiency versus PEmax. Because the mutations in xPE are not known to reduce Cas9 activity (*21*, *23*, *27*), we reasoned that this reduced efficiency might be related to nick repositioning. Previous work showed that pegRNA 3’ ends are unstable, decreasing efficiency by eliminating binding to nicked DNA 3’ ends (*11*). We hypothesized that prime editors with repositioned nicks might be particularly susceptible to pegRNA instability due to reduced overlap between their shortened nicked DNA 3’ ends and pegRNAs. To evaluate this, we swapped our error-suppressing Cas9n from xPE (R221K-K848A-H982A-N1317R) into a recent efficiency-boosting prime editing architecture (PE7) that stabilizes pegRNAs using the La poly-U RNA vPE to PE7 across six loci (*CXCR4*, *EMX1*, *GFP*, *MYC*, *STAT1*, and *TGFB1*) in HEK293T cells using pegRNA-only editing. Compared to PE7, vPE reduced indel rates 8.6-fold (range 1.3 – 16-fold), with similar efficiency, and increased the edit:indel ratio 8.2-fold (range 1.0 – 15-fold) (Fig. 2B). Comparing vPE and PE7 to xPE and PEmax for the same edits (fig. S5, A and B), we observed substantially increased editing efficiency (Fig. 2C). For PE7 versus PEmax, both composed of the same Cas9n (R221K-N394K), the increase in efficiency was 2.7-fold (range 1.4 – 5.3-fold). Yet for vPE versus xPE, the gain in efficiency was 3.2-fold (range 1.5 – 6.5-fold), suggesting xPE was somewhat more restrained by pegRNA instability than PEmax. This increased efficiency coincided with similarly increased indel error rates for PE7 but not for vPE (Fig. 2C). Correspondingly, PE7 featured an edit:indel ratio of 138:1, slightly better than PEmax at 91:1, whereas vPE increased the edit:indel ratio to 465:1 (Fig. 2D).

**Figure 2.**
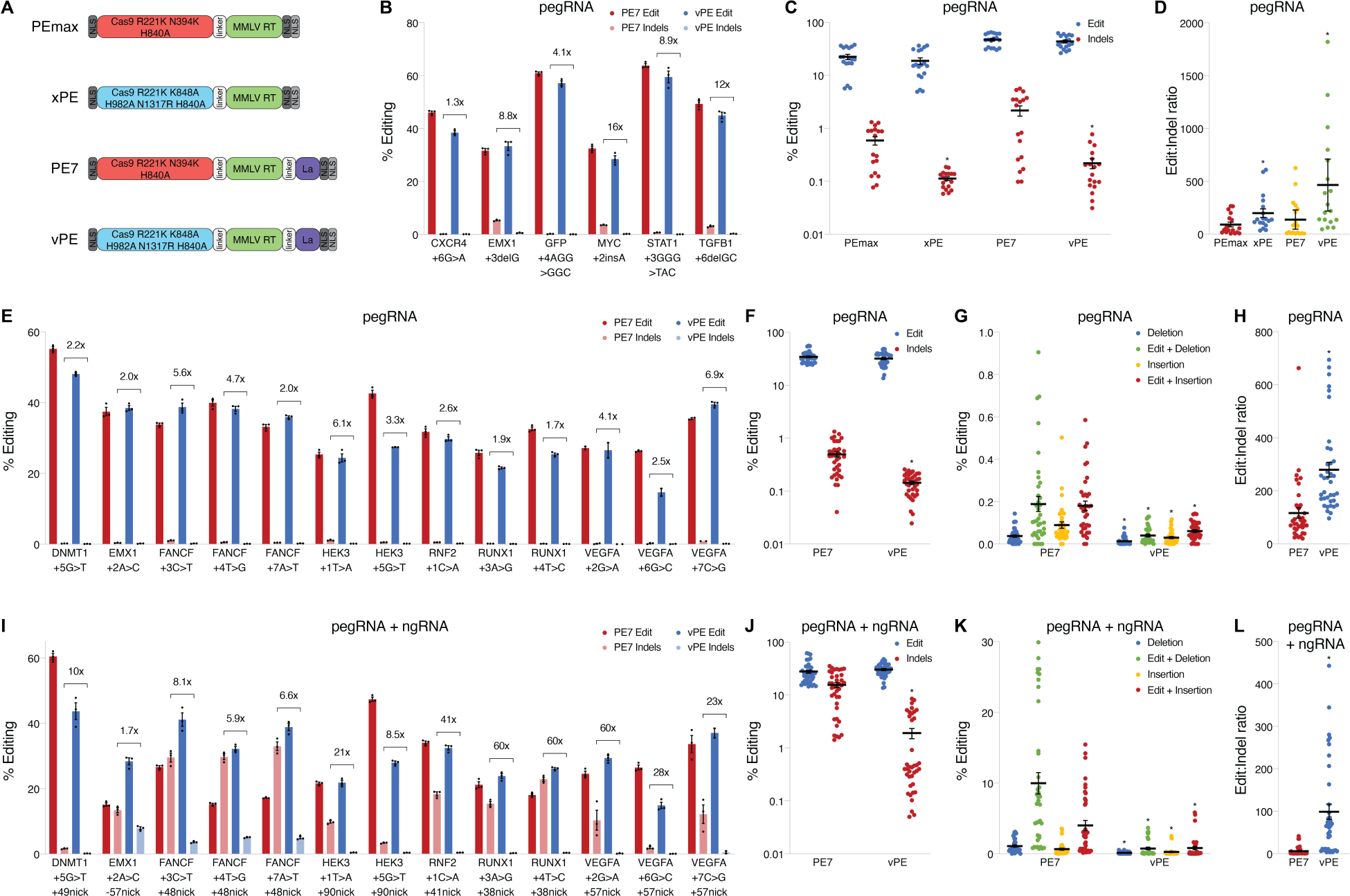
Error-suppression synergizes with pegRNA-stabilizing architecture for efficient editing. (**A**) Architectures of different prime editors used in this study, highlighting the origins of vPE components. (**B**) Edit and indel frequencies comparing vPE to PE7 using pegRNA-only prime editing at several loci, with fold-reductions in indel rates marked. (**C**) Means of edit and indel frequencies comparing PEmax, xPE, PE7, and vPE using pegRNA-only prime editing at several loci (from **B** and **fig. S5A**), with each point representing an individual edit and replicate. (**D**) Means of edit:indel ratios comparing PEmax, xPE, PE7, and vPE using pegRNA-only prime editing at several loci (from **B** and **fig. S5A**), with each point representing an individual edit and replicate. (**E**) Edit and indel frequencies comparing vPE to PE7 using pegRNA-only prime editing at several loci, with fold-reductions in indel rates marked. (**F**) Means of edit and indel frequencies comparing vPE to PE7 using pegRNA-only prime editing at several loci, with each point representing an individual edit and replicate. (**G**) Means of different indel class frequencies comparing vPE to PE7 using pegRNA-only prime editing at several loci, with each point representing an individual edit and replicate. (**H**) Means of edit:indel ratios comparing vPE to PE7 using pegRNA-only prime editing at several loci, with each point representing an individual edit and replicate. (**I**) Edit and indel frequencies comparing vPE to PE7 using pegRNA+ngRNA prime editing at several loci, with fold-reductions in indel rates marked. (**J**) Means of edit and indel frequencies comparing vPE to PE7 using pegRNA+ngRNA prime editing at several loci, with each point representing an individual edit and replicate. (**K**) Means of different indel class frequencies comparing vPE to PE7 using pegRNA+ngRNA prime editing at several loci, with each point representing an individual edit and replicate. (**L**) Means of edit:indel ratios comparing vPE to PE7 using pegRNA+ngRNA prime editing at several loci, with each point representing an individual edit and replicate. * indicates p < 0.05 for comparisons to PEmax in **C** and **D**, and to PE7 in **F**-**H** and **J**-**L**. All data were analyzed by deep sequencing and represent means of *n* = 2-3 independent replicates with standard errors.

To clarify the potential of vPE over earlier prime editors, we edited a larger set of targets for both pegRNA- only and pegRNA+ngRNA editing modes. Since previous optimization of pegRNAs and paired ngRNAs enabled low indel error rates (*2*), we tested these optimized designs to evaluate whether further suppression was achievable (*9*). These edits encompassed all transition and transversion point mutations, spanning +1 to +7 positions, at well-studied loci (*DNMT1*, *EMX1*, *FANCF*, *HEK3/LINC01509*, *RNF2*, *RUNX1*, and *VEGFA*) in HEK293T cells. For pegRNA-only editing, we measured mean efficiencies of 34% (range 25 – 55%) for PE7 and 32% (range 15 – 48%) for vPE (Fig. 2, E and F). For these same edits, we observed mean indel errors of 0.50% (range 0.15 – 1.1%) for PE7 and 0.14% (range 0.039 – 0.23%) for vPE (Fig. 2, E and F). Analyzing indel subtypes revealed that vPE reduced all classes versus PE7 (Fig. 2G). This high efficiency for vPE coupled with significant error suppression led to large increases in edit:indel ratios versus PE7 (Fig. 2H). For pegRNA+ngRNA editing at these same loci and edits, we measured mean editing efficiencies of 28% (range 15 – 60%) for PE7 and 31% (range 15 – 44%) for vPE (Fig. 2, I and J). For pegRNA+ngRNA editing we observed higher mean indel error rates of 16% (range 1.6 – 33%) for PE7 and 1.9% (range 0.069 – 7.9%) for vPE (Fig. 2, I and J). We again found that vPE reduced all indel classes when compared with PE7 (Fig. 2K). Since vPE dramatically reduced errors and increased efficiency for pegRNA+ngRNA editing versus PE7, we observed large increases in edit:indel ratios (Fig. 2L). We further analyzed the allelic identities of edited loci to determine whether errors were dominated by any particular sequence (greater than 0.05% of sequenced reads). Whereas PE7 produced several dominant indel-containing alleles for each edit for pegRNA-only or pegRNA+ngRNA editing, vPE resulted in no significant indel-containing alleles (fig. S6, A-D and fig. S7, A-C). Notably, the fold-reductions in errors were largest for edits where PE7 made the most indels. Thus vPE minimized errors, correcting numerous edits that were highly error-prone with earlier prime editors.

## Discussion

We described the surprising discovery that prime editing errors can be greatly suppressed through engineering the CRISPR nuclease. This finding is explained by repositioning of the Cas9-induced nick that generates resection of competing 5’ strands to reduce their competition with edited 3’ new strands (Fig. 3, A and B). We routinely observed several-fold decreases in indels with our engineered prime editors coupled with high editing efficiencies, resulting in significant increases in edit:indel ratios. This insight suggests how other engineered or natural Cas9 variants and orthologs could be tested for alternative break structures (*28*), which may also promote nontarget strand 5’ end resection following cleavage. Our engineering strategy could potentially be applied to related genome editor classes that similarly utilize polymerases to introduce edits, which also produce significant errors (*29*, *30*). Indeed, we propose that modulating DNA break structure to enhance editing and suppress errors is a design paradigm that can yield superior genome editors.

**Figure 3.**
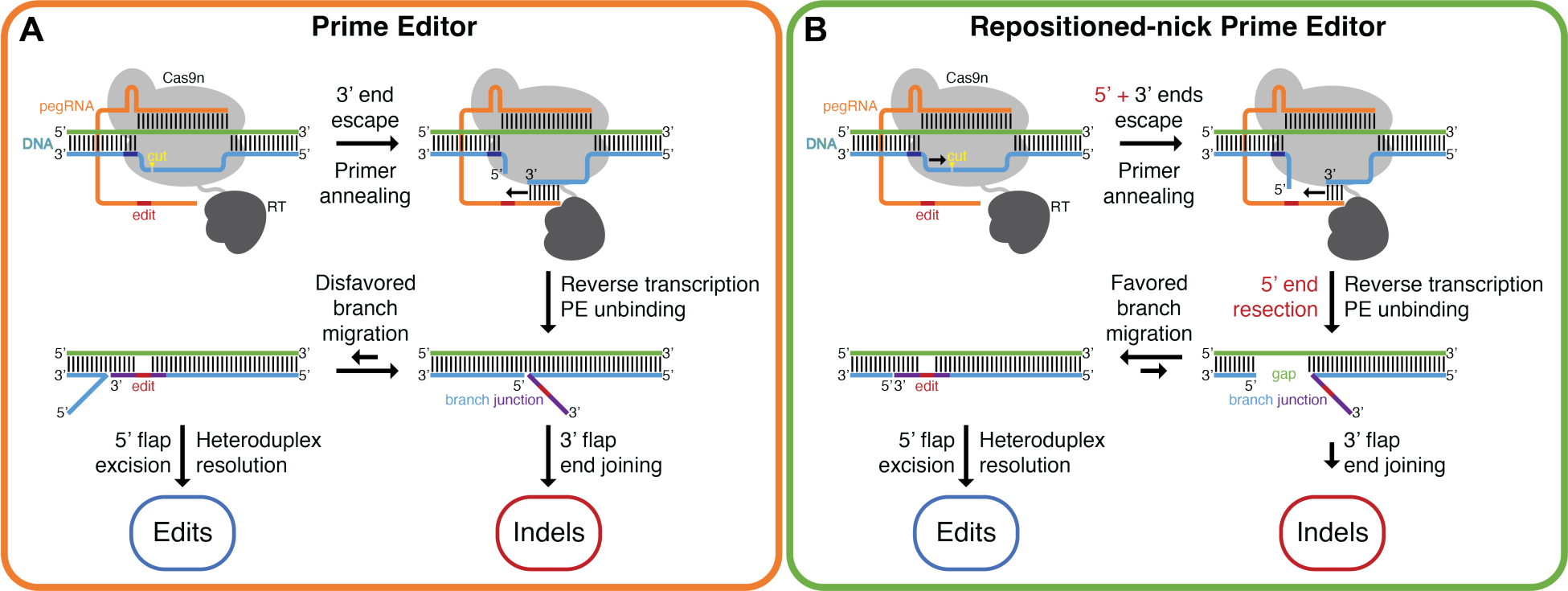
Model for how engineered prime editors suppress error formation. (**A**) Previous prime editors produce edited 3’ new strands that are disfavored in displacing competing 5’ strands, leading to a structure that promotes indel errors. (**B**) Engineered prime editors with repositioned nicks enable resection of competing 5’ strands, eliminating the barrier to edited 3’ new strand binding and favoring a structure that produces the intended edit.

Uncertainty in editing outcomes is a major concern as errors might propagate with harmful consequences, for example in multiplexed editing, gene drives, molecular recording, and gene therapy. The unpredictability of errors is a significant design challenge as efficiency must be maximized while indel errors are minimized, necessitating extensive pegRNA and ngRNA optimization (*31–33*). The engineered prime editors described here appear to eliminate one of these constraints by suppressing errors to minimal levels. Intriguingly, we observed that pegRNA-only editing with vPE reduces error rates to near-uniformly low levels, in contrast to a large range of error rates for PE7. We speculate that variations in local sequences and cellular factors controlling relative annealing rates of the competing 3’ and 5’ strands influence the edit:indel ratio. This is supported by our observation that vPE consistently suppresses error rates to low levels, regardless of how frequent those errors were for PE7. Sequence differences may also explain why different edits at the same locus can yield very different editing efficiencies for vPE versus PE7. Because these adjacent edits utilize nearly identical pegRNAs, this observation could be explained by vPE being sensitive to sequence-specific misfolding that is known to occur for gRNAs (*34*). This suggests that although vPE may simplify pegRNA design, some optimization to maximize efficiency will likely be necessary.

The design of prime editors that produce minimal errors shows that protein engineering can address a key challenge for precise genome editing. This approach is minimally invasive to cells, requiring no manipulation of cellular states, modulation of DNA repair processes, or addition of exogenous factors. Use of these engineered prime editors is straightforward, enabling their facile substitution into existing and future genome editing applications. The prime editors described here exhibit a uniquely high level of editing precision at many loci and could potentially form the basis for a range of advanced tools and applications.

## Acknowledgements

The authors thank David R. Liu for helpful suggestions and critical reading of the manuscript. We thank Allen Jiang, Dig Bijay Mahat, Francisco J. Sanchez-Rivera, and Jacob Witten for helpful discussions.

## Author Contributions

V.P.C., P.A.S., and R.L. conceived or designed the work; V.P.C., P.A.S., and R.L. acquired, analyzed, or interpreted the data; V.P.C., P.A.S., and R.L. drafted the work or substantively revised it.

## Data and Materials Availability

Raw next-generation sequencing data are to be made available on NCBI SRA. Plasmids are to be deposited on Addgene.

## Supplementary Materials

Materials and Methods Figs. S1 to S7

Tables S1 to S3 Sequences

## References

1. A. V Anzalone, L. W. Koblan, D. R. Liu, Genome editing with CRISPR–Cas nucleases, base editors, transposases and prime editors. Nat. Biotechnol. 38, 824–844 (2020).

2. A. V Anzalone et al., Search-and-replace genome editing without double-strand breaks or donor DNA. Nature. 576, 149–157 (2019).

3. J. Choi et al., Precise genomic deletions using paired prime editing. Nat. Biotechnol. 40, 218–226 (2022).

4. Q. Lin et al., High-efficiency prime editing with optimized, paired pegRNAs in plants. Nat. Biotechnol. 39, 923–927 (2021).

5. A. V Anzalone et al., Programmable deletion, replacement, integration and inversion of large DNA sequences with twin prime editing. Nat. Biotechnol. 40, 731–740 (2022).

6. J. Wang et al., Efficient targeted insertion of large DNA fragments without DNA donors. Nat. Methods. 19, 331–340 (2022).

7. M. T. N. Yarnall et al., Drag-and-drop genome insertion of large sequences without double-strand DNA cleavage using CRISPR-directed integrases. Nat. Biotechnol. 41, 500–512 (2023).

8. C. Zheng et al., Template-jumping prime editing enables large insertion and exon rewriting in vivo. Nat. Commun. 14, 3369 (2023).

9. P. J. Chen et al., Enhanced prime editing systems by manipulating cellular determinants of editing outcomes. Cell. 184, 5635–5652.e29 (2021).

10. J. Ferreira da Silva et al., Prime editing efficiency and fidelity are enhanced in the absence of mismatch repair. Nat. Commun. 13, 760 (2022).

11. J. W. Nelson et al., Engineered pegRNAs improve prime editing efficiency. Nat. Biotechnol. 40, 402–410 (2022).

12. X. Li et al., Highly efficient prime editing by introducing same-sense mutations in pegRNA or stabilizing its structure. Nat. Commun. 13, 1669 (2022).

13. J. Yan et al., Improving prime editing with an endogenous small RNA-binding protein. Nature. 628, 639–647 (2024).

14. 14. B. Liu et al., A split prime editor with untethered reverse transcriptase and circular RNA template. Nat. Biotechnol. 40, 1388–1393 (2022).

15. 15. J. Grünewald et al., Engineered CRISPR prime editors with compact, untethered reverse transcriptases. Nat. Biotechnol. 41, 337–343 (2023).

16. 16. J. L. Doman et al., Phage-assisted evolution and protein engineering yield compact, efficient prime editors. Cell. 186, 3983-4002.e26 (2023).

17. M. Fiumara et al., Genotoxic effects of base and prime editing in human hematopoietic stem cells.Nat. Biotechnol. (2023).

18. J. Lee et al., Prime editing with genuine Cas9 nickases minimizes unwanted indels. Nat. Commun. 14, 1786 (2023).

19. C. D. Richardson, G. J. Ray, M. A. DeWitt, G. L. Curie, J. E. Corn, Enhancing homology-directed genome editing by catalytically active and inactive CRISPR-Cas9 using asymmetric donor DNA. Nat. Biotechnol. 34, 339 (2016).

20. Y. Wang et al., Real-time observation of Cas9 postcatalytic domain motions. Proc. Natl. Acad. Sci. 118, e2010650118 (2021).

21. V. P. Chauhan, P. A. Sharp, R. Langer, Altered DNA repair pathway engagement by engineered CRISPR-Cas9 nucleases. Proc. Natl. Acad. Sci. U. S. A. 120 (2023).

22. B. P. Kleinstiver et al., High-fidelity CRISPR–Cas9 nucleases with no detectable genome-wide off-target effects. Nature. 529, 490–495 (2016).

23. I. M. Slaymaker et al., Rationally engineered Cas9 nucleases with improved specificity. Science. 351, 84–88 (2016).

24. B. P. Kleinstiver et al., Engineered CRISPR-Cas9 nucleases with altered PAM specificities. Nature (2015).

25. H. Nishimasu et al., Engineered CRISPR-Cas9 nuclease with expanded targeting space. Science. (2018).

26. R. T. Walton, K. A. Christie, M. N. Whittaker, B. P. Kleinstiver, Unconstrained genome targeting with near-PAMless engineered CRISPR-Cas9 variants. Science. 368, 290–296 (2020).

27. J. M. Spencer, X. Zhang, Deep mutational scanning of S. pyogenes Cas9 reveals important functional domains. Sci. Rep. 7, 16836 (2017).

28. G. Gasiunas et al., A catalogue of biochemically diverse CRISPR-Cas9 orthologs. Nat. Commun. 11, 5512 (2020).

29. B. Liu et al., Targeted genome editing with a DNA-dependent DNA polymerase and exogenous DNA-containing templates. Nat. Biotechnol. (2023).

30. J. F. da Silva et al., *bioRxiv*, doi:10.1101/2023.09.12.557440.

31. J. Y. Hsu et al., PrimeDesign software for rapid and simplified design of prime editing guide RNAs. Nat. Commun. 12, 1034 (2021).

32. R. D. Chow, J. S. Chen, J. Shen, S. Chen, A web tool for the design of prime-editing guide RNAs. *Nat*. Biomed. Eng. 5, 190–194 (2021).

33. H. K. Kim et al., Predicting the efficiency of prime editing guide RNAs in human cells. Nat. Biotechnol. 39, 198–206 (2021).

34. S. B. Thyme, L. Akhmetova, T. G. Montague, E. Valen, A. F. Schier, Internal guide RNA interactions interfere with Cas9-mediated cleavage. Nat. Commun. 7, 11750 (2016).

35. K. Clement et al., CRISPResso2 provides accurate and rapid genome editing sequence analysis. Nat. Biotechnol. 37, 224–226 (2019).

