## Supplementary Materials for "Engineered prime editors with minimal genomic errors"

**Materials and Methods**

Mammalian cell culture

All mammalian cell cultures were maintained in a 37°C incubator at 5% $CO_2$. HEK293T human embryonic kidney cells were maintained in Dulbecco's Modified Eagle's Medium with high glucose, sodium pyruvate, and GlutaMAX (DMEM; ThermoFisher, 10569) supplemented with 10% Fetal Bovine Serum (FBS; ThermoFisher, 10438), and 100 U/mL Penicillin-Streptomycin (ThermoFisher, 15140). HEK293T cells without or with a GFP transgene insertion were used as previously described (*21*).

Mutagenesis and cloning

PE2 and PEmax prime editors were obtained from pCMV-PE2 and pCMV-PEmax, and a cloning backbone for pegRNA expression was obtained from pU6-pegRNA-GG-acceptor, which were gifts from David Liu. PE7 prime editor was obtained from Lenti-PE7-P2A-Puro, which was a gift from Francisco J. Sanchez-Rivera. PE was created by restriction cloning of Cas9n (H840A) from pCMV-PE2 into pCMV-PEmax using NotI and SacI digestion. PE mutagenesis was performed using PCR-driven splicing by overlap extension using primers listed in Table S1. Briefly, one fragment was amplified by PCR from PE using the pe-FWD or pe-mid-FWD and *mutant*-BOT primers and a second fragment was amplified using the *mutant*-TOP and pe-mid-REV or pe-rt-REV primers for each mutant. Each pair of fragments was then spliced by overlap extension PCR using the pe-FWD and pe-mid-REV or pe-mid-FWD and pe-rt-REV primers to create a PE gene fragment with a single residue mutation. These PE gene fragments were then each cloned back into PE using unique NotI, SacI, and BamHI restriction sites to replace the PE sequence with the mutant sequence. Additional mutants (double-, triple-, and quadruple-mutants) were made iteratively starting from these single-mutant plasmids. PE7 and vPE were created by restriction cloning of a fragment containing La amplified from Lenti-PE7-P2A-Puro into pCMV-PEmax, for PE7, or into xPE, for vPE, using BamHI and BshTI digestion. The pegRNA oligos, listed in Table S2, were cloned into pU6-pegRNA-GG-acceptor by Golden Gate cloning with Eco31I digestion.

A custom gRNA cloning backbone vector was created by PCR amplification from pX330 using the gRNA-scaffold-NheI-FWD and gRNA-scaffold-EcoRI-REV primers and restriction cloning into pUC19 (ThermoFisher) using NheI and EcoRI digestion. The nicking gRNA spacer sequence oligos, listed in Table S2, were phosphorylated with T4 polynucleotide kinase (NEB) and cloned into gRNA cloning backbone by Golden Gate cloning with BpiI digestion.

Primers were synthesized by IDT. Restriction enzymes were obtained from ThermoFisher. T7 DNA ligase was obtained from NEB. Plasmids were transformed into Stbl3 chemically competent *E. Coli* (ThermoFisher). Sequences for the PEmax, PE, pPE, xPE, PE7, and vPE vectors are presented in the Sequences section.

Structure analysis

Crystal structures of Cas9 with substrate DNA bound (5F9R) or without substrate DNA bound (4ZT0) were analyzed using PyMol (Schrödinger).

Cell transfection

Cells were seeded in the maintenance medium without Pen-Strep into 48-well plates at 50,000 cells/well. Transfections of HEK293T with prime editing vectors were carried out 24 hrs after seeding using 238 ng PE expression vector, 57 ng pegRNA expression vector, and 72 ng nicking gRNA expression vector (for pegRNA+ngRNA editing) formulated with 0.74-0.92 $\mu$L (equal volume / DNA) Lipofectamine 2000 at a total volume of 29.5-36.7 $\mu$L (equal DNA concentration) in OptiMEM I (ThermoFisher) per well. For sequencing assays, genomic DNA was extracted 72 hrs after transfection using QuickExtract (Epicentre).

High-throughput sequencing

The targeted loci were amplified from extracted genomic DNA by PCR using Herculase II polymerase (Agilent). The PCR primers included Illumina sequencing handles as well as replicate-specific barcodes. These PCR products were then tagged with sample-specific barcodes and sequenced on an Illumina MiSeq. Primers, listed in Table S3, were synthesized by IDT.

Genome editing analysis

To measure editing outcomes, the high-throughput sequencing data were analyzed using CRISPResso2 (*35*). Data for prime editing experiments were processed using the 'prime editing' mode in CRISPResso2 by including sequence values for the parameters 'prime_editing_pegRNA_spacer_seq,' 'prime_editing_pegRNA_extension_seq,' 'prime_editing_pegRNA_scaffold_seq,' and 'prime_editing_nicking_guide_seq' (for pegRNA+ngRNA modes). Editing window parameters ‘prime_editing_pegRNA_extension_quantification_window_size’ and ‘w’ were set to 5. The ‘ignore_substitutions’ option was used to account for small sequence variations that occur due to PCR and sequencing errors. Intended edit rates were quantified as the fraction of reads marked as prime edited out of total sequencing reads. Indel rates were quantified as the fraction of reads marked as indels out of total sequencing reads. Frequencies of specific indel sizes were quantified as the fraction of reads containing these sizes out of all indel reads or out of total sequencing reads, as noted, and were averaged over three independent replicates. Mean indel sizes were calculated as the mean of the absolute values of indel sizes weighted by their indel fractions. Depletion of specific indel sizes was quantified as the fractional reduction in the frequency of that indel size, comparing different editors. Plots of insertion and deletion positions were produced from data generated in CRISPResso2 and averaged over three independent replicates.

Plots of editing outcome alleles were processed using the standard mode in CRISPResso2. The editing window parameter ‘w’ was set to 30. The plot size parameter ‘plot_window_size’ was set to 30, the minimum allele frequency parameter ‘min_frequency_alleles_around_cut_to_plot’ was set to 0.05, and the allele number parameter ‘max_rows_alleles_around_cut_to_plot’ was set to 30. Accordingly, the top 30 alleles were displayed regardless of frequency.

<u>DNA nick position and resection analysis</u>

To measure DNA nick position and resection for Cas9 variants, editing outcomes for dual-gRNA cutting of genomic DNA were re-analyzed from previously published data (*21*). In this data set, HEK293T cells were edited with pairs of gRNAs targeting the EMX1 locus. The gRNA pairs were complementary to the same strand at each locus and were expected to make cuts 84 bp apart, resulting in junctions. The loci were amplified and sequenced by high-throughput sequencing. The high-throughput sequencing data were analyzed using CRISPResso2 with the expected junction as a reference sequence. To assess DNA nick position, sequencing reads aligned to the junction reference were analyzed for insertion sequences perfectly matching the sequences flanking the expected gRNA cut sites. The positions of these matching sequences at the two gRNA sites were used to determine cut positions leading to each read. The most frequent cut position with a frequency greater than 1% of reads was taken as the nick position. To assess DNA resection, sequencing reads aligned to the junction reference were analyzed for deletion sequences. The lengths of deletions to the PAM-proximal side of the junction were taken as the lengths of resection. The resection length was quantified as the median length of these resection products. The resection frequency was quantified as the fraction of reads containing deletions on the PAM-proximal side of the junction out of all reads containing the junction sequence.

### Edit notation

Edits were denoted based on the position where the edit begins relative to expected gRNA nick position for wild-type Cas9, denoting position +1 as 3bp upstream of the first PAM position. Substitution edits were noted using a ‘>’ mark, deletions were noted by a ‘del’ mark, and insertions were noted by an ‘ins’ mark. The base identities of the strand containing the gRNA spacer sequence were used in all cases.

### Statistical analysis

Specific statistical comparisons are indicated in the figure legends. Error bars indicate the standard error for independent replicates as noted. Significance where noted was assessed using unpaired, two-tailed Student’s t-tests. Figures and analysis were produced using Prism software.

### Supplementary Figures

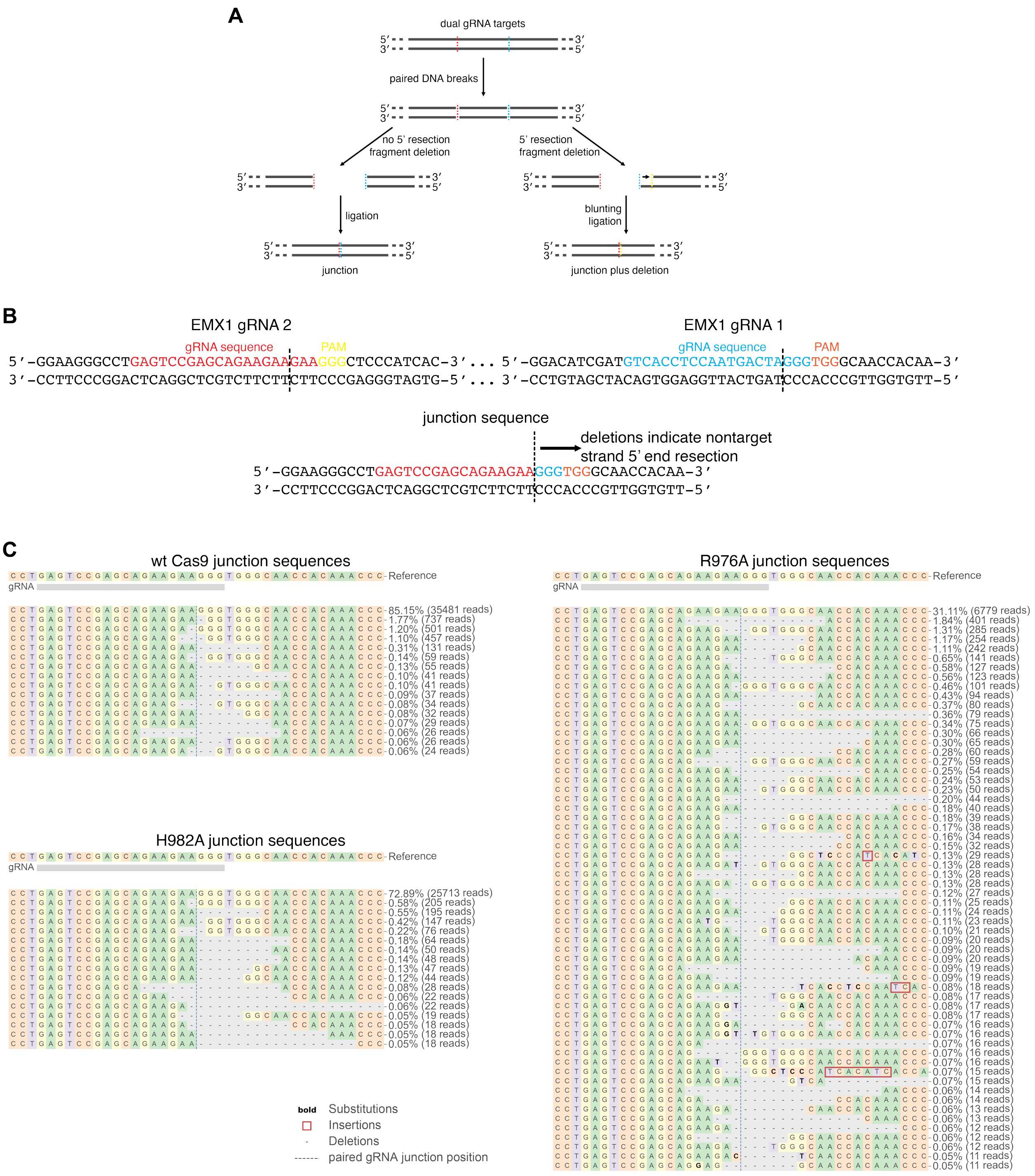


**Figure S1. Assay for measuring nontarget strand 5' end resection.** (**A**) Assay schematic, where paired gRNAs induce DNA breaks that lead to perfect junctions for non-resected ends and deletions in the junction products indicate resection of the 5' DNA ends. The length of each deletion to the PAM-proximal side of the junction indicates the degree of resection, and the frequencies of these deletions indicate the frequency of resection. (**B**) Design of gRNA pairs and expected sequence of the junction product, with the direction of resection-associated deletions indicated. (**C**) Rates of the top genomic sequences resulting from editing with different Cas9 variants at the junction. Only reads with the reference sequence or deletions are depicted. The most frequent allelic sequences above 0.05% of reads are displayed along with sequencing reads and percentages out of all reads. Data were analyzed by deep sequencing and represent a single replicate.

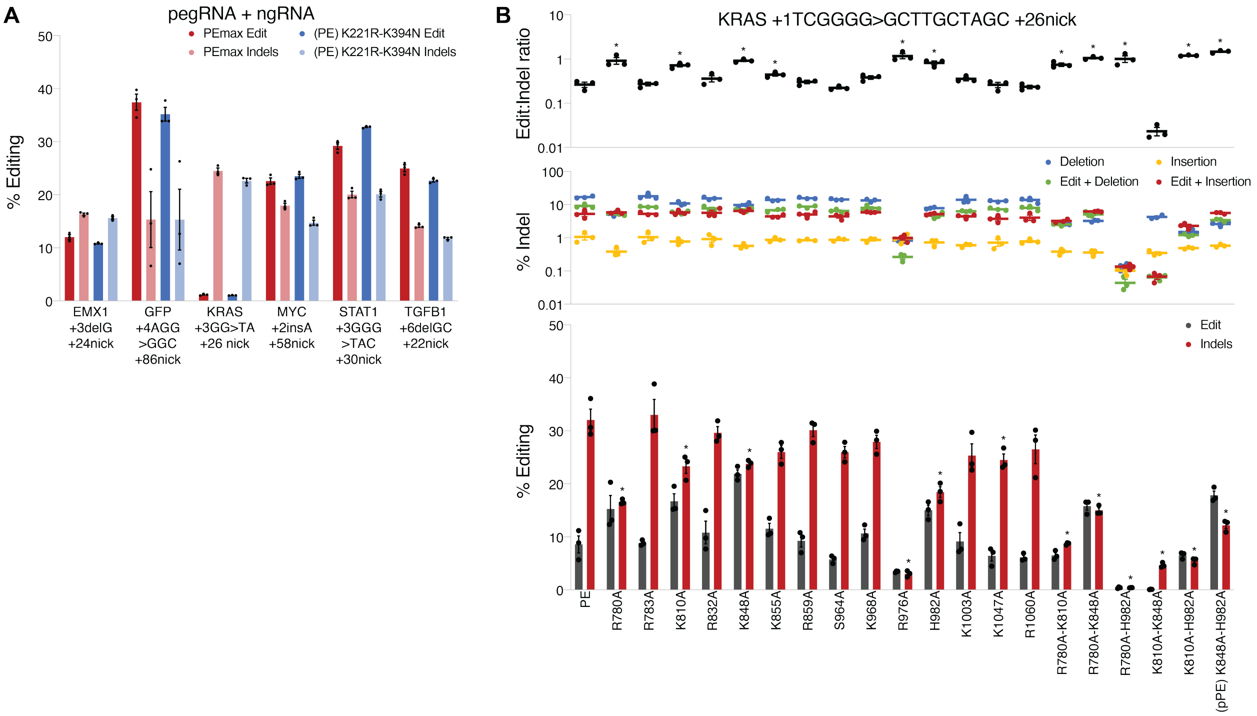


**Figure S2. Effects of prime editor mutations on editing precision.** (**A**) Edit and indel frequencies comparing PE to PEmax using pegRNA+ngRNA prime editing modes at several loci. (**B**) Screen of engineered PE variants to suppress indels with quantification of edit and indel frequencies (bottom), indel classes (middle), and edit:indel ratios (top). * indicates $p < 0.05$ for comparisons to PEmax in **A**, and to PE in **B**. All data were analyzed by deep sequencing and represent means of $n = 3$ independent replicates with standard errors.

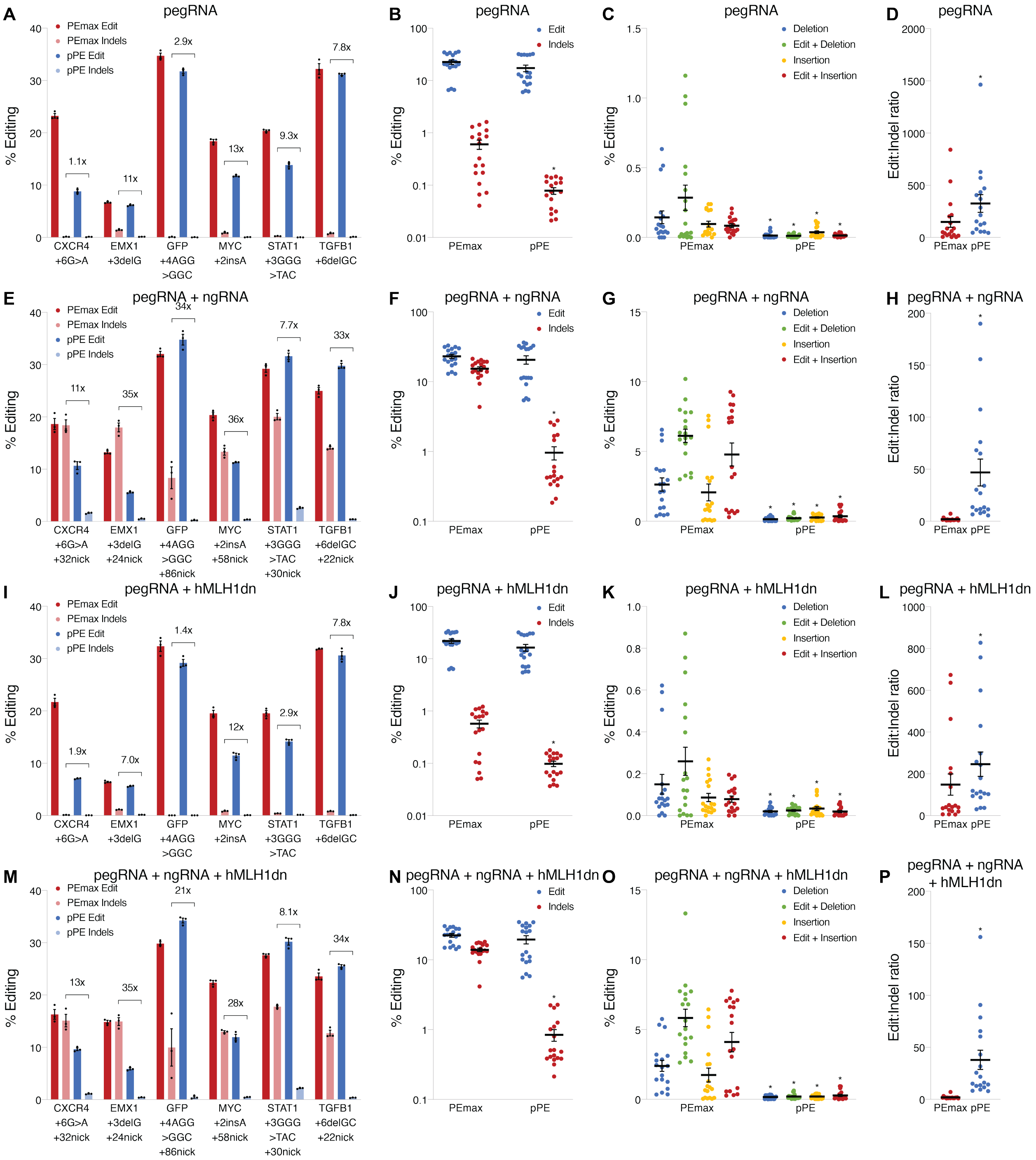


**Figure S3. Comparison of edit and indel generation for pPE and PEmax.** (**A**) Edit and indel frequencies comparing pPE to PEmax using pegRNA-only prime editing at several loci, with fold-reductions in indel rates marked. (**B**) Means of edit and indel frequencies comparing pPE to PEmax using pegRNA-only prime editing at several loci, with each point representing an individual edit and replicate. (**C**) Means of different indel class frequencies comparing pPE to PEmax using pegRNA-only prime editing at several loci, with each point representing an individual edit and replicate. (**D**) Means of edit:indel ratios comparing pPE to PEmax using pegRNA-only prime editing at several loci, with each point representing an individual edit and replicate. (**E**) Edit and indel frequencies comparing pPE to PEmax using pegRNA+ngRNA prime editing at several loci, with fold-reductions in indel rates marked. (**F**) Means of edit and indel frequencies comparing pPE to PEmax using pegRNA+ngRNA prime editing at several loci, with each point representing an individual edit and replicate. (**G**) Means of different indel class frequencies comparing pPE to PEmax using pegRNA+ngRNA prime editing at several loci, with each point representing an individual edit and replicate. (**H**) Means of edit:indel ratios comparing pPE to PEmax using pegRNA+ngRNA prime editing at several loci, with each point representing an individual edit and replicate. (**I**) Edit and indel frequencies comparing pPE to PEmax using pegRNA-only prime editing with inhibition of mismatch repair at several loci, with fold-reductions in indel rates marked. (**J**) Means of edit and indel frequencies comparing pPE to PEmax using pegRNA-only prime editing with inhibition of mismatch repair at several loci, with each point representing an individual edit and replicate. (**K**) Means of different indel class frequencies comparing pPE to PEmax using pegRNA-only prime editing with inhibition of mismatch repair at several loci, with each point representing an individual edit and replicate. (**L**) Means of edit:indel ratios comparing pPE to PEmax using pegRNA-only prime editing with inhibition of mismatch repair at several loci, with each point representing an individual edit and replicate. (**M**) Edit and indel frequencies comparing pPE to PEmax using pegRNA+ngRNA prime editing with inhibition of mismatch repair at several loci, with fold-reductions in indel rates marked. (**N**) Means of edit and indel frequencies comparing pPE to PEmax using pegRNA+ngRNA prime editing with inhibition of mismatch repair at several loci, with each point representing an individual edit and replicate. (**O**) Means of different indel class frequencies comparing pPE to PEmax using pegRNA+ngRNA prime editing with inhibition of mismatch repair at several loci, with each point representing an individual edit and replicate. (**P**) Means of edit:indel ratios comparing pPE to PEmax using pegRNA+ngRNA prime editing with inhibition of mismatch repair at several loci, with each point representing an individual edit and replicate. * indicates $p < 0.05$ for comparisons to PEmax. All data were analyzed by deep sequencing and represent means of $n = 3$ independent replicates with standard errors.

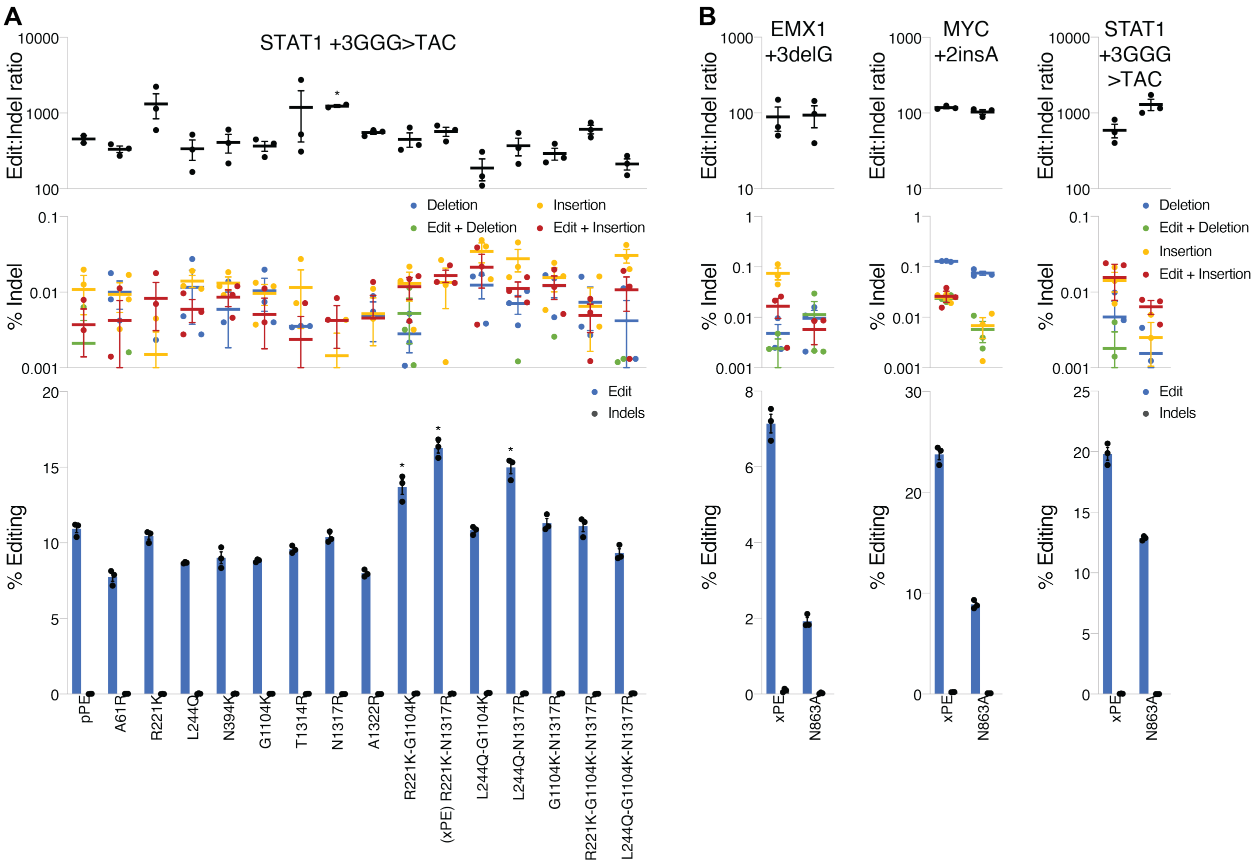


**Figure S4. Further engineering of pPE to rescue lost editing efficiency.** (**A**) Screen of engineered pPE variants to enhance efficiency with quantification of edit and indel frequencies (bottom), indel classes (middle), and edit:indel ratios (top). (**B**) Comparison of xPE and its N863A variant with quantification of edit and indel frequencies (bottom), indel classes (middle), and edit:indel ratios (top). * indicates $p < 0.05$ for comparisons to pPE. All data were analyzed by deep sequencing and represent means of $n = 3$ independent replicates with standard errors.

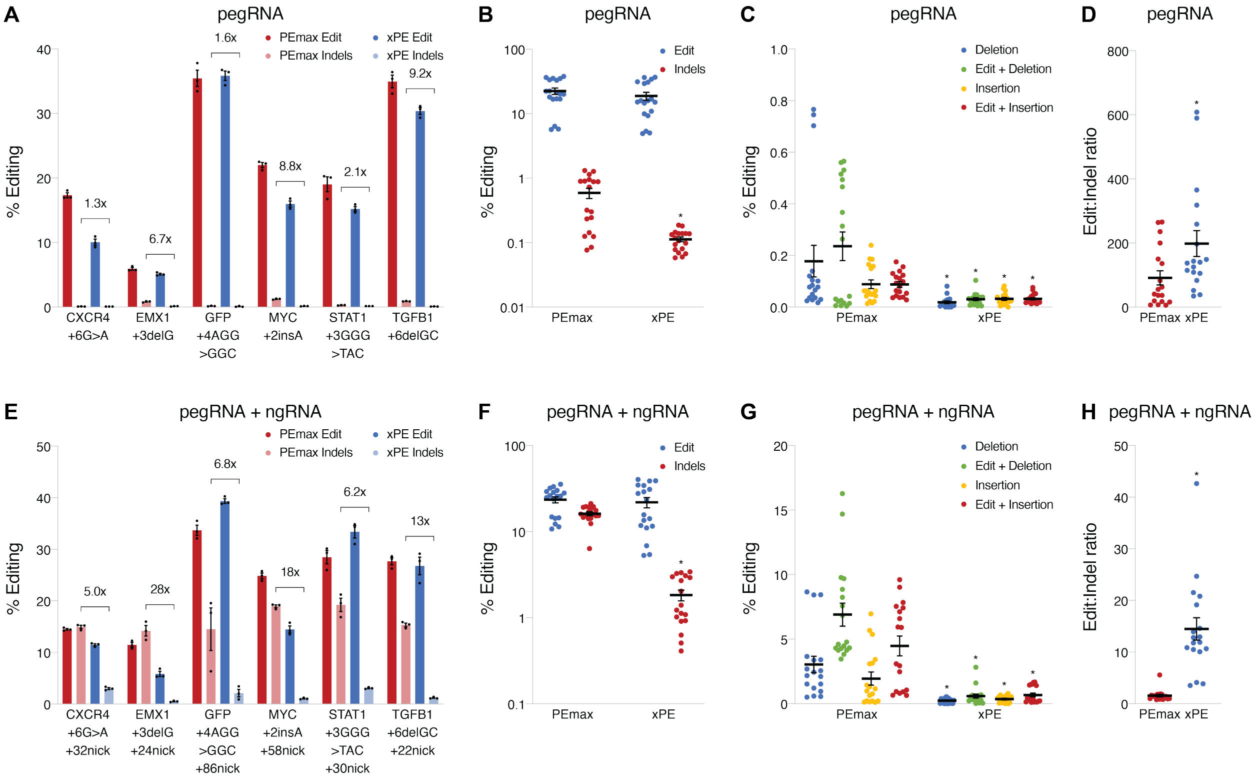


**Figure S5. Comparison of edit and indel generation for xPE and PEmax.** (**A**) Edit and indel frequencies comparing xPE to PEmax using pegRNA-only prime editing at several loci, with fold-reductions in indel rates marked. (**B**) Means of edit and indel frequencies comparing xPE to PEmax using pegRNA-only prime editing at several loci, with each point representing an individual edit and replicate. (**C**) Means of different indel class frequencies comparing xPE to PEmax using pegRNA-only prime editing at several loci, with each point representing an individual edit and replicate. (**D**) Means of edit:indel ratios comparing xPE to PEmax using pegRNA-only prime editing at several loci, with each point representing an individual edit and replicate. (**E**) Edit and indel frequencies comparing xPE to PEmax using pegRNA+ngRNA prime editing at several loci, with fold-reductions in indel rates marked. (**F**) Means of edit and indel frequencies comparing xPE to PEmax using pegRNA+ngRNA prime editing at several loci, with each point representing an individual edit and replicate. (**G**) Means of different indel class frequencies comparing xPE to PEmax using pegRNA+ngRNA prime editing at several loci, with each point representing an individual edit and replicate. (**H**) Means of edit:indel ratios comparing xPE to PEmax using pegRNA+ngRNA prime editing at several loci, with each point representing an individual edit and replicate. * indicates $p < 0.05$ for comparisons to PEmax. All data were analyzed by deep sequencing and represent means of $n = 3$ independent replicates with standard errors.

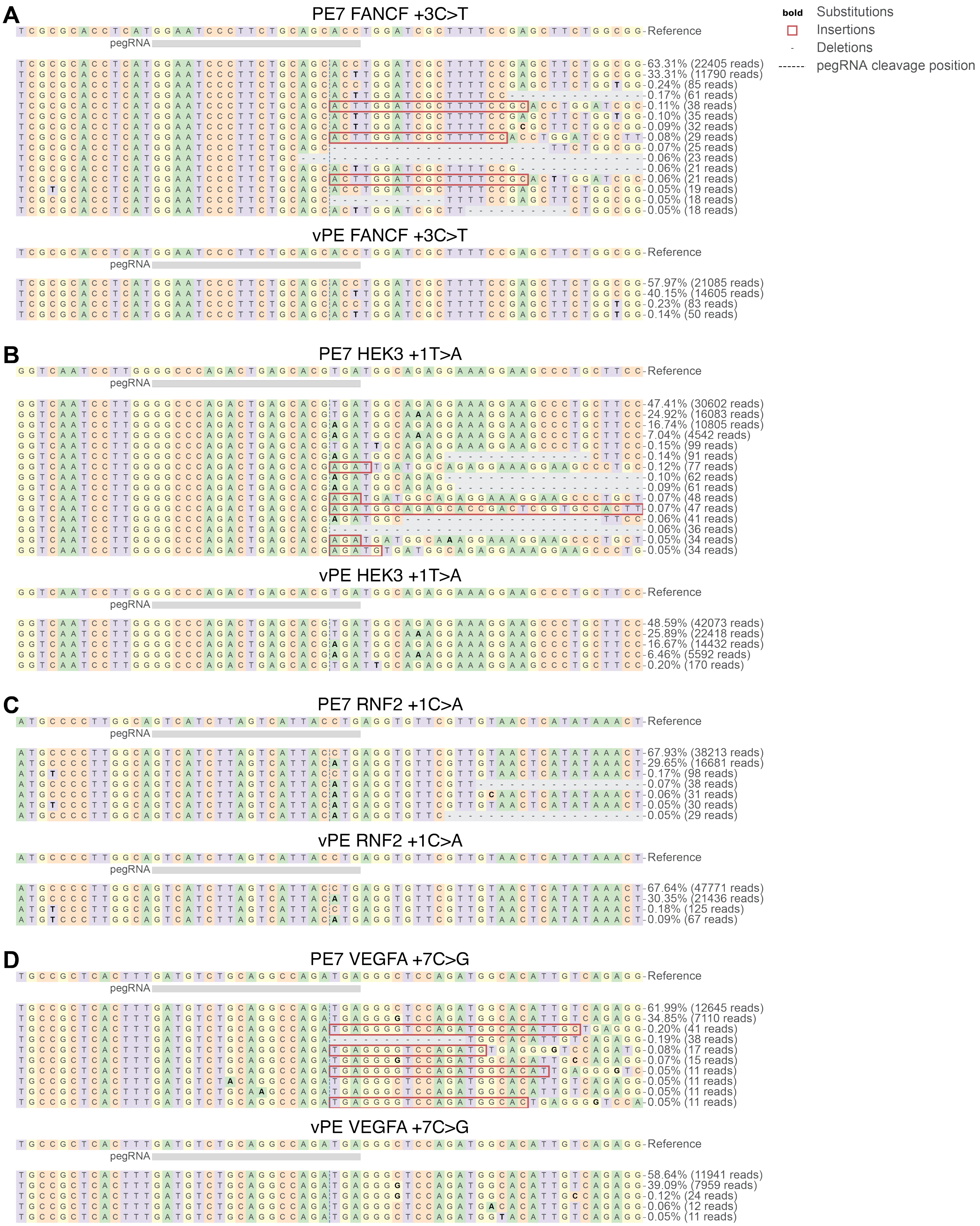


**Figure S6. Sequencing reads showing pegRNA-only prime editing products for PE7 and vPE.** (**A**-**D**) Rates of the top genomic sequences resulting from editing with PE7 or vPE at the (**A**) FANCF, (**B**) HEK3,

(**C**) RNF2, and (**D**) VEGFA loci. Substitutions, insertions, and deletions are depicted. The most frequent allelic sequences above 0.05% of reads are displayed along with sequencing reads and percentages out of all reads. Data were analyzed by deep sequencing and represent a single replicate.

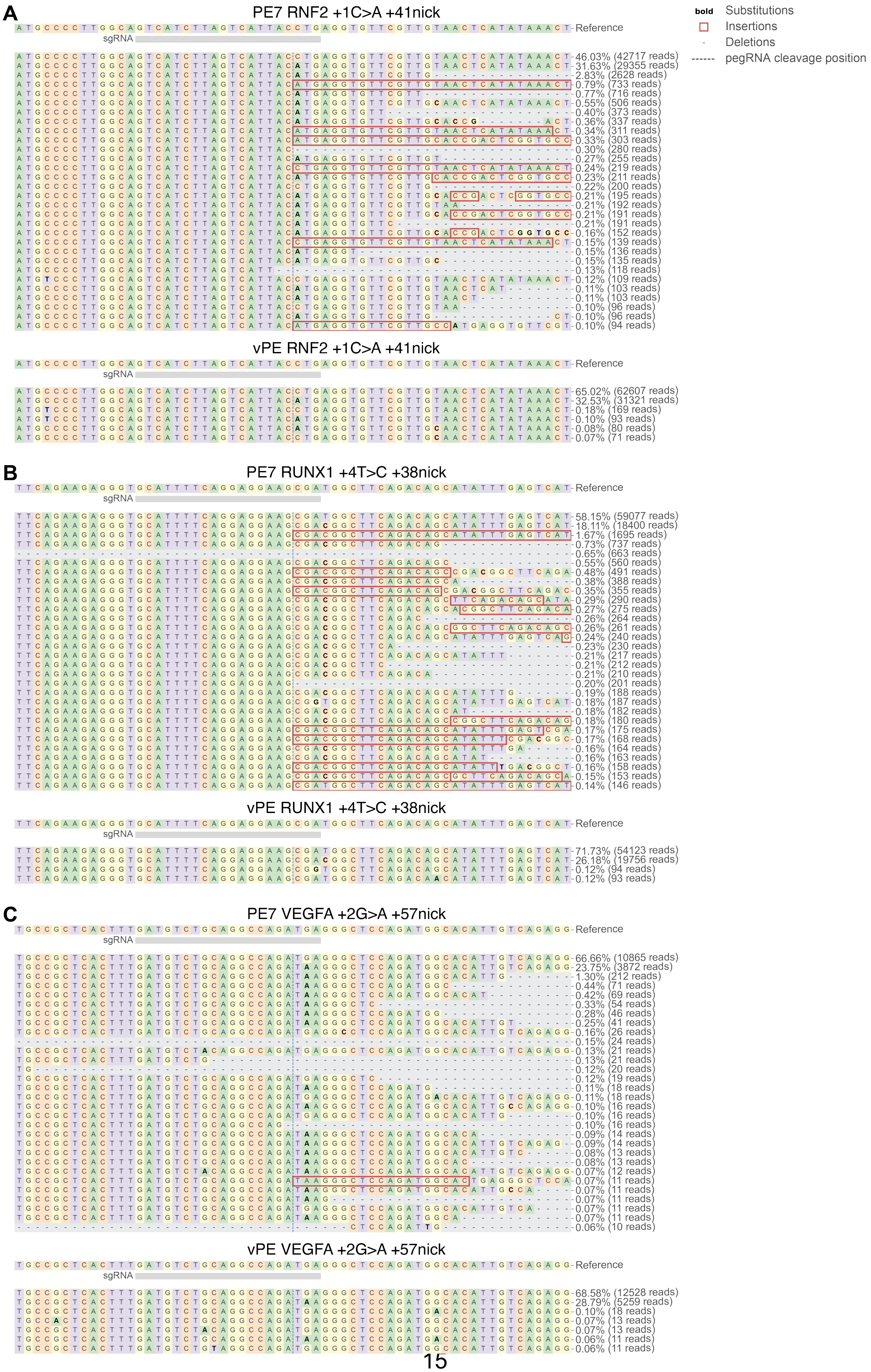
A
PE7 RNF2 +1C>A +41nick
vPE RNF2 +1C>A +41nick
B
PE7 RUNX1 +4T>C +38nick
vPE RUNX1 +4T>C +38nick
C
PE7 VEGFA +2G>A +57nick
vPE VEGFA +2G>A +57nick
Reference
sgRNA
bold Substitutions
Insertions
Deletions
pegRNA cleavage position

**Figure S7. Sequencing reads showing pegRNA+ngRNA prime editing products for PE7 and vPE.** (**A**-**C**) Rates of the top genomic sequences resulting from editing with PE7 or vPE at the (**A**) RNF2, (**B**) RUNX1, and (**C**) VEGFA loci. Substitutions, insertions, and deletions are depicted. The most frequent allelic sequences above 0.05% of reads are displayed along with sequencing reads and percentages out of all reads. Data were analyzed by deep sequencing and represent a single replicate.

**Supplementary Tables**

| Name | Sequence (5' - 3') |
|---|---|
| pe-FWD | GCTAGAGATCCGCGGCCGCTAATAC |
| pe-mid-REV | CACTTTCACGAGCTCGTCCACCAC |
| pe-mid-FWD | GTGGTGGACGAGCTCGTGAAAGTG |
| pe-rt-REV | CTGGGTGCTGGATCCGGAAATCTG |
| R780A-BOT | CCCTCTTCGATCCGCTTCATGGCCTCGCGGCTGTTCTTCTGTCC |
| R780A-TOP | GGACAGAAGAACAGCCGCGAGGCCATGAAGCGGATCGAAGAGGG |
| R783A-BOT | GCTCTTTGATGCCCTCTTCGATGGCCTTCATTCTCTCGCGGCTGTTC |
| R783A-TOP | GAACAGCCGCGAGAGAATGAAGGCCATCGAAGAGGGCATCAAAGAGC |
| K810A-BOT | TCTGCAGGTAGTACAGGTACAGGGCCTCGTTCTGCAGCTGGGTGTTT |
| K810A-TOP | AAACACCCAGCTGCAGAACGAGGCCCTGTACCTGTACTACCTGCAGA |
| R832A-BOT | CCACATCGTAGTCGGACAGGGCGTTGATGTCCAGTTCCTGGTCC |
| R832A-TOP | GGACCAGGAACTGGACATCAACGCCCTGTCCGACTACGATGTGG |
| K848A-BOT | CCTTGTTGTCGATGGAGTCGTCGGCCAGAAAGCTCTGAGGCACGATA |
| K848A-TOP | TATCGTGCCTCAGAGCTTTCTGGCCGACGACTCCATCGACAACAAGG |
| K855A-BOT | TTGTCGCTTCTGGTCAGCACGGCGTTGTCGATGGAGTCGTCCT |
| K855A-TOP | AGGACGACTCCATCGACAACGCCGTGCTGACCAGAAGCGACAA |
| S964A-BOT | AAATCCTTCCGGAAATCGGCCACCAGCTTGGACTTCAG |
| S964A-TOP | CTGAAGTCCAAGCTGGTGGCCGATTTCCGGAAGGATTT |
| K968A-BOT | CGCACTTTGTAAAACTGGAAATCGGCCCGGAAATCGGACACCAGCTTGG |
| K968A-TOP | CCAAGCTGGTGTCCGATTTCCGGGCCGATTTCCAGTTTTACAAAGTGCG |
| R976A-BOT | GCGTGGTGGTAGTTGTTGATCTCGGCCACTTTGTAAAACTGGAAATCCT |
| R976A-TOP | AGGATTTCCAGTTTTACAAAGTGGCCGAGATCAACAACTACCACCACGC |
| H982A-BOT | GTAGGCGTCGTGGGCGTGGGCGTAGTTGTTGATCTCGCG |
| H982A-TOP | CGCGAGATCAACAACTACGCCCACGCCCACGACGCCTAC |
| K1003A-BOT | CGTACACGAACTCGCTTTCCAGGGCAGGGTACTTTTTGATCAGGGCG |
| K1003A-TOP | CGCCCTGATCAAAAAGTACCCTGCCCTGGAAAGCGAGTTCGTGTACG |
| K1047A-BOT | GCCGTTGGCCAGGGTAATCTCGGTGGCGAAAAAGTTCATGATGTTGCTG |
| K1047A-TOP | CAGCAACATCATGAACTTTTTCGCCACCGAGATTACCCTGGCCAACGGC |
| R1060A-BOT | CCGTTTGTCTCGATCAGAGGGGCCTTCCGGATCTCGCCGTTGG |
| R1060A-TOP | CCAACGGCGAGATCCGGAAGGCCCCTCTGATCGAGACAAACGG |
| A61R-BOT | GTTCTCTTCAGCCGGGTCCGCTCGGCTGTTTCGCC |
| A61R-TOP | GGCGAAACAGCCGAGCGGACCCGGCTGAAGAGAAC |
| R221K-BOT | GCGATCAGATTTTCCAGCTTTCTGCTCTTGCTCAGTCTG |
| R221K-TOP | CAGACTGAGCAAGAGCAGAAAGCTGGAAAATCTGATCGC |
| L244Q-BOT | GTCAGGCCCAGGCTCTGGGCAATCAGGTTTC |
| L244Q-TOP | GAAACCTGATTGCCCAGAGCCTGGGCCTGAC |
| N394K-BOT | CGCAGCAGGTCCTCTCTCTTCAGCTTCACGAGCAG |

| | |
|---|---|
| N394K-TOP | CTGCTCGTGAAGCTGAAGAGAGAGGACCTGCTGCG |
| G1104K-BOT | GGATAGACTCTTTGCTGAACTTGCCTGTCTGCACCTCGGTC |
| G1104K-TOP | GACCGAGGTGCAGACAGGCAAGTTCAGCAAAGAGTCTATCC |
| T1314R-BOT | CAGGGGCTCCCAGATTGGTCAGCCGAAACAGGTGGATGATATTCTCG |
| T1314R-TOP | CGAGAATATCATCCACCTGTTTCGGCTGACCAATCTGGGAGCCCCTG |
| N1317R-BOT | AGGCGGCAGGGGCTCCCAGCCGGGTCAGGGTAAACAGGTGG |
| N1317R-TOP | CCACCTGTTTACCCTGACCCGGCTGGGAGCCCCTGCCGCCT |
| A1322R-BOT | GTGGTGTCAAAGTACTTGAAGGCCCGAGGGGCTCCCAGATTGGTCAGGG |
| A1322R-TOP | CCCTGACCAATCTGGGAGCCCCTCGGGCCTTCAAGTACTTTGACACCAC |
| N863A-BOT | GTTGTCGCTCTTGCCCCGGGCCTTGTCGCTTCTGGTCAGC |
| N863A-TOP | GCTGACCAGAAGCGACAAGGCCCGGGGCAAGAGCGACAAC |

**Table S1. Oligodeoxynucleotide sequences used for mutagenesis and cloning.**

| Name | Sequence (5' - 3') |
|---|---|
| pegRNA-scaffold-BOT | /5Phos/GCACCGACTCGGTGCCACTTTTTCAAGTTGATAACGGACTAGCCTTATTTTAACTTGCTATTTCTAG |
| pegRNA-scaffold-TOP | /5Phos/AGAGCTAGAAATAGCAAGTTAAAATAAGGCTAGTCCGTTATCAACTTGAAAAAGTGGCACCGAGTCG |
| CXCR4-pegS1-BOT | CTCTAAAACCCTCTTTGTCATCACGCTTC |
| CXCR4-pegS1-TOP | CACCGAAGCGTGATGACAAAGAGGGTTTT |
| CXCR4-+6G>A-pegT-BOT | AAAAAGCGTGATGACAAAGAGGAGATCGGCCACTGACA |
| CXCR4-+6G>A-pegT-TOP | GTGCTGTCAGTGGCCGATCTCCTCTTTGTCATCACGCT |
| CXCR4-nickgRNA1-BOT | AAACGTACTTGTCCGTCATGCTTC |
| CXCR4-nickgRNA1-TOP | CACCGAAGCATGACGGACAAGTAC |
| DNMT1-pegS1-BOT | CTCTAAAACTGTTTCTGGCACCAGGAATC |
| DNMT1-pegS1-TOP | CACCGATTCCTGGTGCCAGAAACAGTTTT |
| DNMT1-+5G>T-pegT-BOT | AAAACCTGGTGCCAGAAACAGTGGTGAC |
| DNMT1-+5G>T-pegT-TOP | GTGCGTCACCACTGTTTCTGGCACCAGG |
| DNMT1-nickgRNA1-BOT | AAACCCTTTATTTTAGCTGAAGGGC |
| DNMT1-nickgRNA1-TOP | CACCGCCCTTCAGCTAAAATAAAGG |
| EMX1-pegS1-BOT | CTCTAAAACCCCTAGTCATTGGAGGTGAC |
| EMX1-pegS1-TOP | CACCGTCACCTCCAATGACTAGGGGTTTT |
| EMX1-+3delG-pegT-BOT | AAAACACCTCCAATGACTAGGTGGGCAACCACAAACCC |
| EMX1-+3delG-pegT-TOP | GTGCGGGTTTGTGGTTGCCCACCTAGTCATTGGAGGTG |
| EMX1-nickgRNA1-BOT | AAACCGAGGGCAGAGTGCTGCTTGC |
| EMX1-nickgRNA1-TOP | CACCGCAAGCAGCACTCTGCCCTCG |
| EMX1-pegS2-BOT | CTCTAAAACTTCTTCTTCTGCTCGGACTC |
| EMX1-pegS2-TOP | CACCGAGTCCGAGCAGAAGAAGAAGTTTT |
| EMX1-+2A>C-pegT-BOT | AAAATCCGAGCAGAAGAAGCAGGGCTCCCATCAC |
| EMX1-+2A>C-pegT-TOP | GTGCGTGATGGGAGCCCTGCTTCTTCTGCTCGGA |
| EMX1-nickgRNA2-BOT | AAACGAGGACAAAGTACAAACGGC |
| EMX1-nickgRNA2-TOP | CACCGCCGTTTGTACTTTGTCCTC |
| FANCF-pegS1-BOT | CTCTAAAACGGTGCTGCAGAAGGGATTCC |
| FANCF-pegS1-TOP | CACCGGAATCCCTTCTGCAGCACCGTTTT |
| FANCF-+3C>T-pegT-BOT | AAAAATCCCTTCTGCAGCACTTGGATCGCTTTTCC |
| FANCF-+3C>T-pegT-TOP | GTGCGGAAAAGCGATCCAAGTGCTGCAGAAGGGAT |
| FANCF-+4T>G-pegT-BOT | AAAAATCCCTTCTGCAGCACCGGGATCGCTTTTCC |
| FANCF-+4T>G-pegT-TOP | GTGCGGAAAAGCGATCCCGGTGCTGCAGAAGGGAT |
| FANCF-+7A>T-pegT-BOT | AAAAATCCCTTCTGCAGCACCTGGTTCGCTTTTCC |
| FANCF-+7A>T-pegT-TOP | GTGCGGAAAAGCGAACCAGGTGCTGCAGAAGGGAT |
| FANCF-nickgRNA1-BOT | AAACACGTCAGCACCTGGGACCCC |
| FANCF-nickgRNA1-TOP | CACCGGGGTCCCAGGTGCTGACGT |
| GFP-pegS1-BOT | CTCTAAAACACGGCGTGCAGTGCTTCAGC |
| GFP-pegS1-TOP | CACCGCTGAAGCACTGCACGCCGTGTTTT |

| | |
|---|---|
| GFP-+4AGG>GGC-pegT-BOT | AAAAGCACTGCACGCCGTGGCTCAGGGTGGTCACGA |
| GFP-+4AGG>GGC-pegT-TOP | GTGCTCGTGACCACCCTGAGCCACGGCGTGCAGTGC |
| GFP-nickgRNA1-BOT | AAACTAGGTGGCATCGCCCTCGCC |
| GFP-nickgRNA1-TOP | CACCGGCGAGGGCGATGCCACCTA |
| HEK3-pegS1-BOT | CTCTAAAACTCACGTGCTCAGTCTGGGCC |
| HEK3-pegS1-TOP | CACCGGCCCAGACTGAGCACGTGAGTTTT |
| HEK3-+1T>A-pegT-BOT | AAAACAGACTGAGCACGAGATGGCAGA |
| HEK3-+1T>A-pegT-TOP | GTGCTCTGCCATCTCGTGCTCAGTCTG |
| HEK3-+5G>T-pegT-BOT | AAAACAGACTGAGCACGTGATTGCAGA |
| HEK3-+5G>T-pegT-TOP | GTGCTCTGCAATCACGTGCTCAGTCTG |
| HEK3-nickgRNA1-BOT | AAACGCACCGGGATACTGGTTGAC |
| HEK3-nickgRNA1-TOP | CACCGTCAACCAGTATCCCGGTGC |
| KRAS-pegS1-BOT | CTCTAAAACCGATTATTATCAGCCTCAGC |
| KRAS-pegS1-TOP | CACCGCTGAGGCTGATAATAATCGGTTTT |
| KRAS-+1sub6-pegT-BOT | AAAATGAGGCTGATAATAAGCTTGCTAGCCGGCGATCAGACAGCCCCGGT |
| KRAS-+1sub6-pegT-TOP | GTGCACCGGGGCTGTCTGATCGCCGGCTAGCAAGCTTATTATCAGCCTCA |
| KRAS-+3GG>TA-pegT-BOT | AAAAAGGCTGATAATAATCTAGGCGGCGATCAGACA |
| KRAS-+3GG>TA-pegT-TOP | GTGCTGTCTGATCGCCGCCTAGATTATTATCAGCCT |
| KRAS-nickgRNA1-BOT | AAACCGGTGTGGGAAATCGTCCGC |
| KRAS-nickgRNA1-TOP | CACCGCGGACGATTTCCCACACCG |
| MYC-pegS1-BOT | CTCTAAAACGCGTCGGGAGAGTCGCGTCC |
| MYC-pegS1-TOP | CACCGGACGCGACTCTCCCGACGCGTTTT |
| MYC-+2insA-pegT-BOT | AAAAACGCGACTCTCCCGACAGCGGGGAGGCTATTCTGC |
| MYC-+2insA-pegT-TOP | GTGCGCAGAATAGCCTCCCCGCTGTCGGGAGAGTCGCGT |
| MYC-nickgRNA1-BOT | AAACGCTTCTCTGAAAGGCTCTCC |
| MYC-nickgRNA1-TOP | CACCGGAGAGCCTTTCAGAGAAGC |
| RNF2-pegS1-BOT | CTCTAAAACCAGGTAATGACTAAGATGAC |
| RNF2-pegS1-TOP | CACCGTCATCTTAGTCATTACCTGGTTTT |
| RNF2-+1C>A-pegT-BOT | AAAACATCTTAGTCATTACATGAGGTGTTCGTT |
| RNF2-+1C>A-pegT-TOP | GTGCAACGAACACCTCATGTAATGACTAAGATG |
| RNF2-nickgRNA1-BOT | AAACATGTTTTGCTTAATGGTTGAC |
| RNF2-nickgRNA1-TOP | CACCGTCAACCATTAAGCAAAACAT |
| RUNX1-pegS1-BOT | CTCTAAAACTCGCTTCCTCCTGAAAATGC |
| RUNX1-pegS1-TOP | CACCGCATTTTCAGGAGGAAGCGAGTTTT |
| RUNX1-+3A>G-pegT-BOT | AAAAATTTTCAGGAGGAAGCGGTGGCTTCAGACA |
| RUNX1-+3A>G-pegT-TOP | GTGCTGTCTGAAGCCACCGCTTCCTCCTGAAAAT |
| RUNX1-+4T>C-pegT-BOT | AAAAATTTTCAGGAGGAAGCGACGGCTTCAGACA |

| | |
|---|---|
| RUNX1-+4T>C-pegT-TOP | GTGCTGTCTGAAGCCGTCGCTTCCTCCTGAAAAT |
| RUNX1-nickgRNA1-BOT | AAACTCGTACCCACAGTGCTTCATC |
| RUNX1-nickgRNA1-TOP | CACCGATGAAGCACTGTGGGTACGA |
| STAT1-pegS1-BOT | CTCTAAAACCCAGCTGCAAGCATGTCATC |
| STAT1-pegS1-TOP | CACCGATGACATGCTTGCAGCTGGGTTTT |
| STAT1-+3GGG>TAC-pegT-BOT | AAAATGACATGCTTGCAGCTGTACGAAACAGGTTGTCCT |
| STAT1-+3GGG>TAC-pegT-TOP | GTGCAGGACAACCTGTTTCGTACAGCTGCAAGCATGTCA |
| STAT1-nickgRNA1-BOT | AAACATCTTTGCTCCTGGCTGTCC |
| STAT1-nickgRNA1-TOP | CACCGGACAGCCAGGAGCAAAGAT |
| TGFB1-pegS1-BOT | CTCTAAAACCTTGGTGGAAGCGCAGGCTC |
| TGFB1-pegS1-TOP | CACCGAGCCTGCGCTTCCACCAAGGTTTT |
| TGFB1-+6delGC-pegT-BOT | AAAAGCCTGCGCTTCCACCAAGGGAGGAGGACCCCGCT |
| TGFB1-+6delGC-pegT-TOP | GTGCAGCGGGGTCCTCCTCCCTTGGTGGAAGCGCAGGC |
| TGFB1-nickgRNA1-BOT | AAACCTGCGCTAAACGCTGGCAGTC |
| TGFB1-nickgRNA1-TOP | CACCGACTGCCAGCGTTTAGCGCAG |
| VEGFA-pegS1-BOT | CTCTAAAACTCATCTGGCCTGCAGACATC |
| VEGFA-pegS1-TOP | CACCGATGTCTGCAGGCCAGATGAGTTTT |
| VEGFA-+2G>A-pegT-BOT | AAAATCTGCAGGCCAGATAAGGGCTCCAGATGGCACATT |
| VEGFA-+2G>A-pegT-TOP | GTGCAATGTGCCATCTGGAGCCCTTATCTGGCCTGCAGA |
| VEGFA-+6G>C-pegT-BOT | AAAATCTGCAGGCCAGATGAGGCCTCCAGATGGCACATT |
| VEGFA-+6G>C-pegT-TOP | GTGCAATGTGCCATCTGGAGGCCTCATCTGGCCTGCAGA |
| VEGFA-+7C>G-pegT-BOT | AAAATCTGCAGGCCAGATGAGGGGTCCAGATGGCACATT |
| VEGFA-+7C>G-pegT-TOP | GTGCAATGTGCCATCTGGACCCCTCATCTGGCCTGCAGA |
| VEGFA-nickgRNA1-BOT | AAACGCCCTGGGCTCTCTGTACATC |
| VEGFA-nickgRNA1-TOP | CACCGATGTACAGAGAGCCCAGGGC |

**Table S2. Oligodeoxynucleotide sequences used for pegRNA and ngRNA cloning.**

| Name | Sequence (5' - 3') |
|---|---|
| CXCR4-seq-r1-FWD | ACACTCTTTCCCTACACGACGCTCTTCCGATCTNCGTTGGTCATGGGTTACCAGAAGA |
| CXCR4-seq-r2-FWD | ACACTCTTTCCCTACACGACGCTCTTCCGATCTNACGTGGTCATGGGTTACCAGAAGA |
| CXCR4-seq-r3-FWD | ACACTCTTTCCCTACACGACGCTCTTCCGATCTNGTATGGTCATGGGTTACCAGAAGA |
| CXCR4-seq-REV | GACTGGAGTTCAGACGTGTGCTCTTCCGATCTGACTGATGAAGGCCAGGATG |
| DNMT1-seq-r1-FWD | ACACTCTTTCCCTACACGACGCTCTTCCGATCTNATCACGTTAATGTTTCCTGATGGTCC |
| DNMT1-seq-r2-FWD | ACACTCTTTCCCTACACGACGCTCTTCCGATCTNCGAACGTTAATGTTTCCTGATGGTCC |
| DNMT1-seq-r3-FWD | ACACTCTTTCCCTACACGACGCTCTTCCGATCTNTAGACGTTAATGTTTCCTGATGGTCC |
| DNMT1-seq-REV | GACTGGAGTTCAGACGTGTGCTCTTCCGATCTCACAACAGCTTCATGTCAGCC |
| EMX1-seq1-r1-FWD | ACACTCTTTCCCTACACGACGCTCTTCCGATCTNGACCCTGAGTCCGAGCAGAAGAA |
| EMX1-seq1-r2-FWD | ACACTCTTTCCCTACACGACGCTCTTCCGATCTNTGACCTGAGTCCGAGCAGAAGAA |
| EMX1-seq1-r3-FWD | ACACTCTTTCCCTACACGACGCTCTTCCGATCTNACTCCTGAGTCCGAGCAGAAGAA |
| EMX1-seq1-REV | GACTGGAGTTCAGACGTGTGCTCTTCCGATCTAGTGGCCAGAGTCCAGCTT |
| EMX1-seq2-r1-FWD | ACACTCTTTCCCTACACGACGCTCTTCCGATCTNATCCAGCTCAGCCTGAGTGTTGA |
| EMX1-seq2-r2-FWD | ACACTCTTTCCCTACACGACGCTCTTCCGATCTNCGACAGCTCAGCCTGAGTGTTGA |
| EMX1-seq2-r3-FWD | ACACTCTTTCCCTACACGACGCTCTTCCGATCTNTAGCAGCTCAGCCTGAGTGTTGA |
| EMX1-seq2-REV | GACTGGAGTTCAGACGTGTGCTCTTCCGATCTCTCGTGGGTTTGTGGTTGC |
| FANCF-seq-r1-FWD | ACACTCTTTCCCTACACGACGCTCTTCCGATCTNATGAGACGCTGGGAGATTGACATGC |
| FANCF-seq-r2-FWD | ACACTCTTTCCCTACACGACGCTCTTCCGATCTNGATAGACGCTGGGAGATTGACATGC |
| FANCF-seq-r3-FWD | ACACTCTTTCCCTACACGACGCTCTTCCGATCTNTGCAGACGCTGGGAGATTGACATGC |
| FANCF-seq-REV | GACTGGAGTTCAGACGTGTGCTCTTCCGATCTCGATGGATGTGGCGCAGGTAG |
| GFP-seq-r1-FWD | ACACTCTTTCCCTACACGACGCTCTTCCGATCTNATGCGTAAACGGCCACAAGTTCAGC |
| GFP-seq-r2-FWD | ACACTCTTTCCCTACACGACGCTCTTCCGATCTNGATCGTAAACGGCCACAAGTTCAGC |
| GFP-seq-r3-FWD | ACACTCTTTCCCTACACGACGCTCTTCCGATCTNTGCCGTAAACGGCCACAAGTTCAGC |
| GFP-seq-REV | GACTGGAGTTCAGACGTGTGCTCTTCCGATCTTAGTTGCCGTCGTCCTTGAAGA |
| HEK3-seq-r1-FWD | ACACTCTTTCCCTACACGACGCTCTTCCGATCTNGACCCCAGCCAAACTTGTCAACC |
| HEK3-seq-r2-FWD | ACACTCTTTCCCTACACGACGCTCTTCCGATCTNTGACCCAGCCAAACTTGTCAACC |

| | |
|---|---|
| HEK3-seq-r3-FWD | ACACTCTTTCCCTACACGACGCTCTTCCGATCTNACTCCCAGCCAAACTTG TCAACC |
| HEK3-seq-REV | GACTGGAGTTCAGACGTGTGCTCTTCCGATCTATGTGGGCTGCCTAGAAA GG |
| KRAS-seq-r1-FWD | ACACTCTTTCCCTACACGACGCTCTTCCGATCTNGACTTGAAAGGGTCTGT CGTGTTTG |
| KRAS-seq-r2-FWD | ACACTCTTTCCCTACACGACGCTCTTCCGATCTNTGATTGAAAGGGTCTGT CGTGTTTG |
| KRAS-seq-r3-FWD | ACACTCTTTCCCTACACGACGCTCTTCCGATCTNACTTTGAAAGGGTCTGT CGTGTTTG |
| KRAS-seq-REV | GACTGGAGTTCAGACGTGTGCTCTTCCGATCTAAACAAGCAGTCACCAAAA GTGG |
| MYC-seq-r1-FWD | ACACTCTTTCCCTACACGACGCTCTTCCGATCTNCGTCACGAAACTTTGCC CATAGCA |
| MYC-seq-r2-FWD | ACACTCTTTCCCTACACGACGCTCTTCCGATCTNACGCACGAAACTTTGCC CATAGCA |
| MYC-seq-r3-FWD | ACACTCTTTCCCTACACGACGCTCTTCCGATCTNGTACACGAAACTTTGCC CATAGCA |
| MYC-seq-REV | GACTGGAGTTCAGACGTGTGCTCTTCCGATCTAAGTGGACTTCGGTGCTT ACC |
| RNF2-seq-r1-FWD | ACACTCTTTCCCTACACGACGCTCTTCCGATCTNCGTAGACCATAGCACTT CCCTTCC |
| RNF2-seq-r2-FWD | ACACTCTTTCCCTACACGACGCTCTTCCGATCTNACGAGACCATAGCACTT CCCTTCC |
| RNF2-seq-r3-FWD | ACACTCTTTCCCTACACGACGCTCTTCCGATCTNGTAAGACCATAGCACTT CCCTTCC |
| RNF2-seq-REV | GACTGGAGTTCAGACGTGTGCTCTTCCGATCTTTAGCCAACATACAGAAGT CAGG |
| RUNX1-seq-r1-FWD | ACACTCTTTCCCTACACGACGCTCTTCCGATCTNGACAGATGTAGGGCTAG AGGGGTG |
| RUNX1-seq-r2-FWD | ACACTCTTTCCCTACACGACGCTCTTCCGATCTNTGAAGATGTAGGGCTAG AGGGGTG |
| RUNX1-seq-r3-FWD | ACACTCTTTCCCTACACGACGCTCTTCCGATCTNACTAGATGTAGGGCTAG AGGGGTG |
| RUNX1-seq-REV | GACTGGAGTTCAGACGTGTGCTCTTCCGATCTTCACAAACAAGACAGGGA ACTG |
| STAT1-seq-r1-FWD | ACACTCTTTCCCTACACGACGCTCTTCCGATCTNATGAAAGTAGTATGCGT GGGCCTC |
| STAT1-seq-r2-FWD | ACACTCTTTCCCTACACGACGCTCTTCCGATCTNGATAAAGTAGTATGCGT GGGCCTC |
| STAT1-seq-r3-FWD | ACACTCTTTCCCTACACGACGCTCTTCCGATCTNTGCAAAGTAGTATGCGT GGGCCTC |
| STAT1-seq-REV | GACTGGAGTTCAGACGTGTGCTCTTCCGATCTGCTCAAAAGCTGGTAAAC CTTCA |
| TGFB1-seq-r1-FWD | ACACTCTTTCCCTACACGACGCTCTTCCGATCTNGACGTGACTCTACAAGA CCGAGGTG |
| TGFB1-seq-r2-FWD | ACACTCTTTCCCTACACGACGCTCTTCCGATCTNTGAGTGACTCTACAAGA CCGAGGTG |
| TGFB1-seq-r3-FWD | ACACTCTTTCCCTACACGACGCTCTTCCGATCTNACTGTGACTCTACAAGA CCGAGGTG |
| TGFB1-seq-REV | GACTGGAGTTCAGACGTGTGCTCTTCCGATCTCCTGAGAGGAACTGGGAC TTTG |
| VEGFA-seq-r1- | ACACTCTTTCCCTACACGACGCTCTTCCGATCTNATGACTTGGTGCCAAAT |

| | |
|---|---|
| FWD | TCTTCTCC |
| VEGFA-seq-r2-FWD | ACACTCTTTCCCTACACGACGCTCTTCCGATCTNGATACTTGGTGCCAAATTCTTCTCC |
| VEGFA-seq-r3-FWD | ACACTCTTTCCCTACACGACGCTCTTCCGATCTNTGCACTTGGTGCCAAATTCTTCTCC |
| VEGFA-seq-REV | GACTGGAGTTCAGACGTGTGCTCTTCCGATCTAAAGAGGGAATGGGCTTTGGA |

**Table S3. Oligodeoxynucleotide sequences used for next-generation sequencing.**

**Sequences**

PEmax, with mutations (versus wild-type Cas9) in red

ATGAAACGGACAGCCGACGGAAGCGAGTTCGAGTCACCAAAGAAGAAGCGGAAAGTCGACAAGAAG
TACAGCATCGGCCTGGACATCGGCACCAACTCTGTGGGCTGGGCCGTGATCACCGACGAGTACAAG
GTGCCCAGCAAGAAATTCAAGGTGCTGGGCAACACCGACCGGCACAGCATCAAGAAGAACCTGATC
GGAGCCCTGCTGTTCGACAGCGGCGAAACAGCCGAGGCCACCCGGCTGAAGAGAACCGCCAGAAG
AAGATACACCAGACGGAAGAACCGGATCTGCTATCTGCAAGAGATCTTCAGCAACGAGATGGCCAA
GGTGGACGACAGCTTCTTCCACAGACTGGAAGAGTCCTTCCTGGTGGAAGAGGATAAGAAGCACGA
GCGGCACCCCATCTTCGGCAACATCGTGGACGAGGTGGCCTACCACGAGAAGTACCCCACCATCTA
CCACCTGAGAAAGAAACTGGTGGACAGCACCGACAAGGCCGACCTGCGGCTGATCTATCTGGCCCT
GGCCCACATGATCAAGTTCCGGGGCCACTTCCTGATCGAGGGCGACCTGAACCCCGACAACAGCGA
CGTGGACAAGCTGTTCATCCAGCTGGTGCAGACCTACAACCAGCTGTTCGAGGAAAACCCCATCAA
CGCCAGCGGCGTGGACGCCAAGGCCATCCTGTCTGCCAGACTGAGCAAGAGCAGAAAGCTGGAAA
ATCTGATCGCCCAGCTGCCCGGCGAGAAGAAGAATGGCCTGTTCGGAAACCTGATTGCCCTGAGCC
TGGGCCTGACCCCCAACTTCAAGAGCAACTTCGACCTGGCCGAGGATGCCAAACTGCAGCTGAGCA
AGGACACCTACGACGACGACCTGGACAACCTGCTGGCCCAGATCGGCGACCAGTACGCCGACCTG
TTTCTGGCCGCCAAGAACCTGTCCGACGCCATCCTGCTGAGCGACATCCTGAGAGTGAACACCGAG
ATCACCAAGGCCCCCCTGAGCGCCTCTATGATCAAGAGATACGACGAGCACCACCAGGACCTGACC
CTGCTGAAAGCTCTCGTGCGGCAGCAGCTGCCTGAGAAGTACAAAGAGATTTTCTTCGACCAGAGC
AAGAACGGCTACGCCGGCTACATTGACGGCGGAGCCAGCCAGGAAGAGTTCTACAAGTTCATCAAG
CCCATCCTGGAAAAGATGGACGGCACCGAGGAACTGCTCGTGAAGCTGAAGAGAGAGGACCTGCT
GCGGAAGCAGCGGACCTTCGACAACGGCAGCATCCCCCACCAGATCCACCTGGGAGAGCTGCACG
CCATTCTGCGGCGGCAGGAAGATTTTTACCCATTCCTGAAGGACAACCGGGAAAAGATCGAGAAGAT
CCTGACCTTCCGCATCCCCTACTACGTGGGCCCTCTGGCCAGGGGAAACAGCAGATTCGCCTGGAT
GACCAGAAAGAGCGAGGAAACCATCACCCCCTGGAACTTCGAGGAAGTGGTGGACAAGGGCGCTT
CCGCCCAGAGCTTCATCGAGCGGATGACCAACTTCGATAAGAACCTGCCCAACGAGAAGGTGCTGC
CCAAGCACAGCCTGCTGTACGAGTACTTCACCGTGTATAACGAGCTGACCAAAGTGAAATACGTGAC
CGAGGGAATGAGAAAGCCCGCCTTCCTGAGCGGCGAGCAGAAAAAGGCCATCGTGGACCTGCTGT
TCAAGACCAACCGGAAAGTGACCGTGAAGCAGCTGAAAGAGGACTACTTCAAGAAAATCGAGTGCTT
CGACTCCGTGGAAATCTCCGGCGTGGAAGATCGGTTCAACGCCTCCCTGGGCACATACCACGATCT
GCTGAAAATTATCAAGGACAAGGACTTCCTGGACAATGAGGAAAACGAGGACATTCTGGAAGATATC
GTGCTGACCCTGACACTGTTTGAGGACAGAGAGATGATCGAGGAACGGCTGAAAACCTATGCCCAC
CTGTTCGACGACAAAGTGATGAAGCAGCTGAAGCGGCGGAGATACACCGGCTGGGGCAGGCTGAG

CCGGAAGCTGATCAACGGCATCCGGGACAAGCAGTCCGGCAAGACAATCCTGGATTTCCTGAAGTC
CGACGGCTTCGCCAACAGAAACTTCATGCAGCTGATCCACGACGACAGCCTGACCTTTAAAGAGGA
CATCCAGAAAGCCCAGGTGTCCGGCCAGGGCGATAGCCTGCACGAGCACATTGCCAATCTGGCCG
GCAGCCCCGCCATTAAGAAGGGCATCCTGCAGACAGTGAAGGTGGTGGACGAGCTCGTGAAAGTG
ATGGGCCGGCACAAGCCCGAGAACATCGTGATCGAAATGGCCAGAGAGAACCAGACCACCCAGAA
GGGACAGAAGAACAGCCGCGAGAGAATGAAGCGGATCGAAGAGGGCATCAAAGAGCTGGGCAGCC
AGATCCTGAAAGAACACCCCGTGGAAAACACCCAGCTGCAGAACGAGAAGCTGTACCTGTACTACCT
GCAGAATGGGCGGGATATGTACGTGGACCAGGAACTGGACATCAACCGGCTGTCCGACTACGATGT
GGACGCTATCGTGCCTCAGAGCTTTCTGAAGGACGACTCCATCGACAACAAGGTGCTGACCAGAAG
CGACAAGAACCGGGGCAAGAGCGACAACGTGCCCTCCGAAGAGGTCGTGAAGAAGATGAAGAACT
ACTGGCGGCAGCTGCTGAACGCCAAGCTGATTACCCAGAGAAAGTTCGACAATCTGACCAAGGCCG
AGAGAGGCGGCCTGAGCGAACTGGATAAGGCCGGCTTCATCAAGAGACAGCTGGTGGAAACCCGG
CAGATCACAAAGCACGTGGCACAGATCCTGGACTCCCGGATGAACACTAAGTACGACGAGAATGAC
AAGCTGATCCGGGAAGTGAAAGTGATCACCCTGAAGTCCAAGCTGGTGTCCGATTTCCGGAAGGAT
TTCCAGTTTTACAAAGTGCGCGAGATCAACAACTACCACCACGCCCACGACGCCTACCTGAACGCCG
TCGTGGGAACCGCCCTGATCAAAAAGTACCCTAAGCTGGAAAGCGAGTTCGTGTACGGCGACTACA
AGGTGTACGACGTGCGGAAGATGATCGCCAAGAGCGAGCAGGAAATCGGCAAGGCTACCGCCAAG
TACTTCTTCTACAGCAACATCATGAACTTTTTCAAGACCGAGATTACCCTGGCCAACGGCGAGATCC
GGAAGCGGCCTCTGATCGAGACAAACGGCGAAACCGGGGAGATCGTGTGGGATAAGGGCCGGGAT
TTTGCCACCGTGCGGAAAGTGCTGAGCATGCCCCAAGTGAATATCGTGAAAAAGACCGAGGTGCAG
ACAGGCGGCTTCAGCAAAGAGTCTATCCTGCCCAAGAGGAACAGCGATAAGCTGATCGCCAGAAAG
AAGGACTGGGACCCTAAGAAGTACGGCGGCTTCGACAGCCCCACCGTGGCCTATTCTGTGCTGGTG
GTGGCCAAAGTGGAAAAGGGCAAGTCCAAGAAACTGAAGAGTGTGAAAGAGCTGCTGGGGATCACC
ATCATGGAAAGAAGCAGCTTCGAGAAGAATCCCATCGACTTTCTGGAAGCCAAGGGCTACAAAGAAG
TGAAAAAGGACCTGATCATCAAGCTGCCTAAGTACTCCCTGTTCGAGCTGGAAAACGGCCGGAAGA
GAATGCTGGCCTCTGCCGGCGAACTGCAGAAGGGAAACGAACTGGCCCTGCCCTCCAAATATGTGA
ACTTCCTGTACCTGGCCAGCCACTATGAGAAGCTGAAGGGCTCCCCCGAGGATAATGAGCAGAAAC
AGCTGTTTGTGGAACAGCACAAGCACTACCTGGACGAGATCATCGAGCAGATCAGCGAGTTCTCCAA
GAGAGTGATCCTGGCCGACGCTAATCTGGACAAAGTGCTGTCCGCCTACAACAAGCACCGGGATAA
GCCCATCAGAGAGCAGGCCGAGAATATCATCCACCTGTTTACCCTGACCAATCTGGGAGCCCCTGC
CGCCTTCAAGTACTTTGACACCACCATCGACCGGAAGAGGTACACCAGCACCAAAGAGGTGCTGGA
CGCCACCCTGATCCACCAGAGCATCACCGGCCTGTACGAGACACGGATCGACCTGTCTCAGCTGGG
AGGTGACTCCGGCGGAAGCTCTGGTGGCAGCAAGCGGACCGCCGACGGCTCTGAATTCGAGAGCC
CTAAGAAGAAAAGAAAGGTGAGCGGAGGCTCTAGCGGCGGAAGCACCCTGAACATTGAAGACGAGT
ATAGACTGCATGAAACAAGCAAGGAACCCGACGTGTCCCTGGGCTCCACCTGGCTGTCCGACTTTC
CCCAGGCCTGGGCCGAGACAGGAGGAATGGGCCTGGCCGTGCGGCAGGCACCCCTGATCATCCCT

CTGAAGGCCACCTCTACACCCGTGAGCATCAAGCAGTACCCTATGTCTCAGGAGGCCAGACTGGGC
ATCAAGCCTCACATCCAGAGGCTGCTGGACCAGGGCATCCTGGTGCCATGCCAGAGCCCCTGGAAC
ACACCACTGCTGCCCGTGAAGAAGCCAGGCACCAATGACTATAGACCCGTGCAGGATCTGAGAGAG
GTGAACAAGAGGGTGGAGGATATCCACCCCACCGTGCCCAACCCTTACAATCTGCTGTCCGGCCTG
CCCCCTTCTCACCAGTGGTATACAGTGCTGGACCTGAAGGATGCCTTCTTTTGTCTGAGACTGCACC
CTACCAGCCAGCCACTGTTCGCCTTTGAGTGGAGGGACCCTGAGATGGGCATCTCTGGCCAGCTGA
CCTGGACACGCCTGCCTCAGGGCTTCAAGAATAGCCCAACACTGTTTAACGAGGCCCTGCACCGCG
ACCTGGCAGATTTCCGGATCCAGCACCCAGATCTGATCCTGCTGCAGTACGTGGACGATCTGCTGC
TGGCCGCCACCAGCGAGCTGGATTGCCAGCAGGGAACACGCGCCCTGCTGCAGACCCTGGGAAAC
CTGGGATATAGGGCATCCGCCAAGAAGGCCCAGATCTGTCAGAAGCAGGTGAAGTACCTGGGCTAT
CTGCTGAAGGAGGGCCAGAGATGGCTGACAGAGGCCAGGAAGGAGACAGTGATGGGCCAGCCAAC
ACCCAAGACCCCAAGACAGCTGAGGGAGTTCCTGGGCAAAGCAGGATTTTGCAGGCTGTTCATCCC
AGGATTCGCAGAGATGGCAGCACCTCTGTACCCACTGACCAAGCCGGGCACCCTGTTTAATTGGGG
CCCTGACCAGCAGAAGGCCTATCAGGAGATCAAGCAGGCCCTGCTGACAGCACCAGCCCTGGGCC
TGCCAGACCTGACCAAGCCTTTCGAGCTGTTTGTGGATGAGAAGCAGGGCTACGCCAAGGGCGTGC
TGACCCAGAAGCTGGGACCATGGAGACGGCCCGTGGCCTATCTGTCCAAGAAGCTGGACCCAGTG
GCAGCAGGATGGCCACCATGCCTGAGGATGGTGGCAGCAATCGCCGTGCTGACAAAGGATGCCGG
CAAGCTGACCATGGGACAGCCACTGGTCATCCTGGCACCACACGCAGTGGAGGCCCTGGTGAAGC
AGCCTCCAGATCGCTGGCTGTCTAACGCCCGGATGACACACTACCAGGCCCTGCTGCTGGACACCG
ATCGCGTGCAGTTTGGCCCTGTGGTGGCCCTGAATCCAGCCACCCTGCTGCCTCTGCCAGAGGAGG
GCCTGCAGCACAACTGTCTGGACATCCTGGCAGAGGCACACGGAACAAGGCCAGACCTGACCGATC
AGCCCCTGCCTGACGCCGATCACACATGGTATACCGATGGAAGCTCCCTGCTGCAGGAGGGCCAGA
GGAAGGCAGGAGCAGCAGTGACCACAGAGACAGAAGTGATCTGGGCCAAGGCCCTGCCAGCAGGC
ACATCCGCCCAGCGGGCCGAGCTGATCGCCCTGACCCAGGCCCTGAAGATGGCCGAGGGCAAGAA
GCTGAACGTGTACACAGACTCCAGATATGCCTTCGCCACCGCACACATCCACGGAGAGATCTACAG
GCGCCGGGGCTGGCTGACCTCTGAGGGCAAGGAGATCAAGAACAAGGATGAGATCCTGGCCCTGC
TGAAGGCCCTGTTTCTGCCCAAGCGGCTGAGCATCATCCACTGTCCTGGACACCAGAAGGGACACT
CCGCCGAGGCAAGGGGCAATCGGATGGCCGACCAGGCCGCCAGAAAGGCTGCTATTACTGAAACT
CCCGACACTTCCACTCTGCTGATTGAAAACTCCTCCCCTTCTGGCGGCTCAAAAAGAACCGCCGACG
GCAGCGAATTCGAGTCTCCCAAGAAGAAGAGGAAAGTCGGCTCTGGCCCTGCCGCTAAGAGAGTGA
AGCTGGAC

PE

ATGAAACGGACAGCCGACGGAAGCGAGTTCGAGTCACCAAAGAAGAAGCGGAAAGTCGACAAGAAG
TACAGCATCGGCCTGGACATCGGCACCAACTCTGTGGGCTGGGCCGTGATCACCGACGAGTACAAG
GTGCCCAGCAAGAAATTCAAGGTGCTGGGCAACACCGACCGGCACAGCATCAAGAAGAACCTGATC
GGAGCCCTGCTGTTCGACAGCGGCGAAACAGCCGAGGCCACCCGGCTGAAGAGAACCGCCAGAAG
AAGATACACCAGACGGAAGAACCGGATCTGCTATCTGCAAGAGATCTTCAGCAACGAGATGGCCAA
GGTGGACGACAGCTTCTTCCACAGACTGGAAGAGTCCTTCCTGGTGGAAGAGGATAAGAAGCACGA
GCGGCACCCCATCTTCGGCAACATCGTGGACGAGGTGGCCTACCACGAGAAGTACCCCACCATCTA
CCACCTGAGAAAGAAACTGGTGGACAGCACCGACAAGGCCGACCTGCGGCTGATCTATCTGGCCCT
GGCCCACATGATCAAGTTCCGGGGCCACTTCCTGATCGAGGGCGACCTGAACCCCGACAACAGCGA
CGTGGACAAGCTGTTCATCCAGCTGGTGCAGACCTACAACCAGCTGTTCGAGGAAAACCCCATCAA
CGCCAGCGGCGTGGACGCCAAGGCCATCCTGTCTGCCAGACTGAGCAAGAGCAGACGGCTGGAAA
ATCTGATCGCCCAGCTGCCCGGCGAGAAGAAGAATGGCCTGTTCGGAAACCTGATTGCCCTGAGCC
TGGGCCTGACCCCCAACTTCAAGAGCAACTTCGACCTGGCCGAGGATGCCAAACTGCAGCTGAGCA
AGGACACCTACGACGACGACCTGGACAACCTGCTGGCCCAGATCGGCGACCAGTACGCCGACCTG
TTTCTGGCCGCCAAGAACCTGTCCGACGCCATCCTGCTGAGCGACATCCTGAGAGTGAACACCGAG
ATCACCAAGGCCCCCCTGAGCGCCTCTATGATCAAGAGATACGACGAGCACCACCAGGACCTGACC
CTGCTGAAAGCTCTCGTGCGGCAGCAGCTGCCTGAGAAGTACAAAGAGATTTTCTTCGACCAGAGC
AAGAACGGCTACGCCGGCTACATTGACGGCGGAGCCAGCCAGGAAGAGTTCTACAAGTTCATCAAG
CCCATCCTGGAAAAGATGGACGGCACCGAGGAACTGCTCGTGAAGCTGAACAGAGAGGACCTGCTG
CGGAAGCAGCGGACCTTCGACAACGGCAGCATCCCCCACCAGATCCACCTGGGAGAGCTGCACGC
CATTCTGCGGCGGCAGGAAGATTTTTACCCATTCCTGAAGGACAACCGGGAAAAGATCGAGAAGATC
CTGACCTTCCGCATCCCCTACTACGTGGGCCCTCTGGCCAGGGGAAACAGCAGATTCGCCTGGATG
ACCAGAAAGAGCGAGGAAACCATCACCCCCTGGAACTTCGAGGAAGTGGTGGACAAGGGCGCTTCC
GCCCAGAGCTTCATCGAGCGGATGACCAACTTCGATAAGAACCTGCCCAACGAGAAGGTGCTGCCC
AAGCACAGCCTGCTGTACGAGTACTTCACCGTGTATAACGAGCTGACCAAAGTGAAATACGTGACCG
AGGGAATGAGAAAGCCCGCCTTCCTGAGCGGCGAGCAGAAAAAGGCCATCGTGGACCTGCTGTTCA
AGACCAACCGGAAAGTGACCGTGAAGCAGCTGAAAGAGGACTACTTCAAGAAAATCGAGTGCTTCG
ACTCCGTGGAAATCTCCGGCGTGGAAGATCGGTTCAACGCCTCCCTGGGCACATACCACGATCTGC
TGAAAATTATCAAGGACAAGGACTTCCTGGACAATGAGGAAAACGAGGACATTCTGGAAGATATCGT
GCTGACCCTGACACTGTTTGAGGACAGAGAGATGATCGAGGAACGGCTGAAAACCTATGCCCACCT
GTTCGACGACAAAGTGATGAAGCAGCTGAAGCGGCGGAGATACACCGGCTGGGGCAGGCTGAGCC
GGAAGCTGATCAACGGCATCCGGGACAAGCAGTCCGGCAAGACAATCCTGGATTTCCTGAAGTCCG
ACGGCTTCGCCAACAGAAACTTCATGCAGCTGATCCACGACGACAGCCTGACCTTTAAAGAGGACAT
CCAGAAAGCCCAGGTGTCCGGCCAGGGCGATAGCCTGCACGAGCACATTGCCAATCTGGCCGGCA

GCCCCGCCATTAAGAAGGGCATCCTGCAGACAGTGAAGGTGGTGGACGAGCTCGTGAAAGTGATG
GGCCGGCACAAGCCCGAGAACATCGTGATCGAAATGGCCAGAGAGAACCAGACCACCCAGAAGGG
ACAGAAGAACAGCCGCGAGAGAATGAAGCGGATCGAAGAGGGCATCAAAGAGCTGGGCAGCCAGA
TCCTGAAAGAACACCCCGTGGAAAACACCCAGCTGCAGAACGAGAAGCTGTACCTGTACTACCTGC
AGAATGGGCGGGATATGTACGTGGACCAGGAACTGGACATCAACCGGCTGTCCGACTACGATGTGG
ACGCTATCGTGCCTCAGAGCTTTCTGAAGGACGACTCCATCGACAACAAGGTGCTGACCAGAAGCG
ACAAGAACCGGGGCAAGAGCGACAACGTGCCCTCCGAAGAGGTCGTGAAGAAGATGAAGAACTACT
GGCGGCAGCTGCTGAACGCCAAGCTGATTACCCAGAGAAAGTTCGACAATCTGACCAAGGCCGAGA
GAGGCGGCCTGAGCGAACTGGATAAGGCCGGCTTCATCAAGAGACAGCTGGTGGAAACCCGGCAG
ATCACAAAGCACGTGGCACAGATCCTGGACTCCCGGATGAACACTAAGTACGACGAGAATGACAAG
CTGATCCGGGAAGTGAAAGTGATCACCCTGAAGTCCAAGCTGGTGTCCGATTTCCGGAAGGATTTCC
AGTTTTACAAAGTGCGCGAGATCAACAACTACCACCACGCCCACGACGCCTACCTGAACGCCGTCG
TGGGAACCGCCCTGATCAAAAAGTACCCTAAGCTGGAAAGCGAGTTCGTGTACGGCGACTACAAGG
TGTACGACGTGCGGAAGATGATCGCCAAGAGCGAGCAGGAAATCGGCAAGGCTACCGCCAAGTACT
TCTTCTACAGCAACATCATGAACTTTTTCAAGACCGAGATTACCCTGGCCAACGGCGAGATCCGGAA
GCGGCCTCTGATCGAGACAAACGGCGAAACCGGGGAGATCGTGTGGGATAAGGGCCGGGATTTTG
CCACCGTGCGGAAAGTGCTGAGCATGCCCCAAGTGAATATCGTGAAAAAGACCGAGGTGCAGACAG
GCGGCTTCAGCAAAGAGTCTATCCTGCCCAAGAGGAACAGCGATAAGCTGATCGCCAGAAAGAAGG
ACTGGGACCCTAAGAAGTACGGCGGCTTCGACAGCCCCACCGTGGCCTATTCTGTGCTGGTGGTGG
CCAAAGTGGAAAAGGGCAAGTCCAAGAAACTGAAGAGTGTGAAAGAGCTGCTGGGGATCACCATCA
TGGAAAGAAGCAGCTTCGAGAAGAATCCCATCGACTTTCTGGAAGCCAAGGGCTACAAAGAAGTGAA
AAAGGACCTGATCATCAAGCTGCCTAAGTACTCCCTGTTCGAGCTGGAAAACGGCCGGAAGAGAAT
GCTGGCCTCTGCCGGCGAACTGCAGAAGGGAAACGAACTGGCCCTGCCCTCCAAATATGTGAACTT
CCTGTACCTGGCCAGCCACTATGAGAAGCTGAAGGGCTCCCCCGAGGATAATGAGCAGAAACAGCT
GTTTGTGGAACAGCACAAGCACTACCTGGACGAGATCATCGAGCAGATCAGCGAGTTCTCCAAGAG
AGTGATCCTGGCCGACGCTAATCTGGACAAAGTGCTGTCCGCCTACAACAAGCACCGGGATAAGCC
CATCAGAGAGCAGGCCGAGAATATCATCCACCTGTTTACCCTGACCAATCTGGGAGCCCCTGCCGC
CTTCAAGTACTTTGACACCACCATCGACCGGAAGAGGTACACCAGCACCAAAGAGGTGCTGGACGC
CACCCTGATCCACCAGAGCATCACCGGCCTGTACGAGACACGGATCGACCTGTCTCAGCTGGGAGG
TGACTCCGGCGGAAGCTCTGGTGGCAGCAAGCGGACCGCCGACGGCTCTGAATTCGAGAGCCCTA
AGAAGAAAAGAAAGGTGAGCGGAGGCTCTAGCGGCGGAAGCACCCTGAACATTGAAGACGAGTATA
GACTGCATGAAACAAGCAAGGAACCCGACGTGTCCCTGGGCTCCACCTGGCTGTCCGACTTTCCCC
AGGCCTGGGCCGAGACAGGAGGAATGGGCCTGGCCGTGCGGCAGGCACCCCTGATCATCCCTCTG
AAGGCCACCTCTACACCCGTGAGCATCAAGCAGTACCCTATGTCTCAGGAGGCCAGACTGGGCATC
AAGCCTCACATCCAGAGGCTGCTGGACCAGGGCATCCTGGTGCCATGCCAGAGCCCCTGGAACACA
CCACTGCTGCCCGTGAAGAAGCCAGGCACCAATGACTATAGACCCGTGCAGGATCTGAGAGAGGTG

AACAAGAGGGTGGAGGATATCCACCCCACCGTGCCCAACCCTTACAATCTGCTGTCCGGCCTGCCC
CCTTCTCACCAGTGGTATACAGTGCTGGACCTGAAGGATGCCTTCTTTTGTCTGAGACTGCACCCTA
CCAGCCAGCCACTGTTCGCCTTTGAGTGGAGGGACCCTGAGATGGGCATCTCTGGCCAGCTGACCT
GGACACGCCTGCCTCAGGGCTTCAAGAATAGCCCAACACTGTTTAACGAGGCCCTGCACCGCGACC
TGGCAGATTTCCGGATCCAGCACCCAGATCTGATCCTGCTGCAGTACGTGGACGATCTGCTGCTGG
CCGCCACCAGCGAGCTGGATTGCCAGCAGGGAACACGCGCCCTGCTGCAGACCCTGGGAAACCTG
GGATATAGGGCATCCGCCAAGAAGGCCCAGATCTGTCAGAAGCAGGTGAAGTACCTGGGCTATCTG
CTGAAGGAGGGCCAGAGATGGCTGACAGAGGCCAGGAAGGAGACAGTGATGGGCCAGCCAACACC
CAAGACCCCAAGACAGCTGAGGGAGTTCCTGGGCAAAGCAGGATTTTGCAGGCTGTTCATCCCAGG
ATTCGCAGAGATGGCAGCACCTCTGTACCCACTGACCAAGCCGGGCACCCTGTTTAATTGGGGCCC
TGACCAGCAGAAGGCCTATCAGGAGATCAAGCAGGCCCTGCTGACAGCACCAGCCCTGGGCCTGC
CAGACCTGACCAAGCCTTTCGAGCTGTTTGTGGATGAGAAGCAGGGCTACGCCAAGGGCGTGCTGA
CCCAGAAGCTGGGACCATGGAGACGGCCCGTGGCCTATCTGTCCAAGAAGCTGGACCCAGTGGCA
GCAGGATGGCCACCATGCCTGAGGATGGTGGCAGCAATCGCCGTGCTGACAAAGGATGCCGGCAA
GCTGACCATGGGACAGCCACTGGTCATCCTGGCACCACACGCAGTGGAGGCCCTGGTGAAGCAGC
CTCCAGATCGCTGGCTGTCTAACGCCCGGATGACACACTACCAGGCCCTGCTGCTGGACACCGATC
GCGTGCAGTTTGGCCCTGTGGTGGCCCTGAATCCAGCCACCCTGCTGCCTCTGCCAGAGGAGGGC
CTGCAGCACAACTGTCTGGACATCCTGGCAGAGGCACACGGAACAAGGCCAGACCTGACCGATCAG
CCCCTGCCTGACGCCGATCACACATGGTATACCGATGGAAGCTCCCTGCTGCAGGAGGGCCAGAG
GAAGGCAGGAGCAGCAGTGACCACAGAGACAGAAGTGATCTGGGCCAAGGCCCTGCCAGCAGGCA
CATCCGCCCAGCGGGCCGAGCTGATCGCCCTGACCCAGGCCCTGAAGATGGCCGAGGGCAAGAAG
CTGAACGTGTACACAGACTCCAGATATGCCTTCGCCACCGCACACATCCACGGAGAGATCTACAGG
CGCCGGGGCTGGCTGACCTCTGAGGGCAAGGAGATCAAGAACAAGGATGAGATCCTGGCCCTGCT
GAAGGCCCTGTTTCTGCCCAAGCGGCTGAGCATCATCCACTGTCCTGGACACCAGAAGGGACACTC
CGCCGAGGCAAGGGGCAATCGGATGGCCGACCAGGCCGCCAGAAAGGCTGCTATTACTGAAACTC
CCGACACTTCCACTCTGCTGATTGAAAACTCCTCCCCTTCTGGCGGCTCAAAAAGAACCGCCGACGG
CAGCGAATTCGAGTCTCCCAAGAAGAAGAGGAAAGTCGGCTCTGGCCCTGCCGCTAAGAGAGTGAA
GCTGGAC

PE$^{K848A-H982A}$ (pPE), with mutations (versus wild-type Cas9) in red:

ATGAAACGGACAGCCGACGGAAGCGAGTTCGAGTCACCAAAGAAGAAGCGGAAAGTCGACAAGAAG
TACAGCATCGGCCTGGACATCGGCACCAACTCTGTGGGCTGGGCCGTGATCACCGACGAGTACAAG
GTGCCCAGCAAGAAATTCAAGGTGCTGGGCAACACCGACCGGCACAGCATCAAGAAGAACCTGATC
GGAGCCCTGCTGTTCGACAGCGGCGAAACAGCCGAGGCCACCCGGCTGAAGAGAACCGCCAGAAG

AAGATACACCAGACGGAAGAACCGGATCTGCTATCTGCAAGAGATCTTCAGCAACGAGATGGCCAA
GGTGGACGACAGCTTCTTCCACAGACTGGAAGAGTCCTTCCTGGTGGAAGAGGATAAGAAGCACGA
GCGGCACCCCATCTTCGGCAACATCGTGGACGAGGTGGCCTACCACGAGAAGTACCCCACCATCTA
CCACCTGAGAAAGAAACTGGTGGACAGCACCGACAAGGCCGACCTGCGGCTGATCTATCTGGCCCT
GGCCCACATGATCAAGTTCCGGGGCCACTTCCTGATCGAGGGCGACCTGAACCCCGACAACAGCGA
CGTGGACAAGCTGTTCATCCAGCTGGTGCAGACCTACAACCAGCTGTTCGAGGAAAACCCCATCAA
CGCCAGCGGCGTGGACGCCAAGGCCATCCTGTCTGCCAGACTGAGCAAGAGCAGACGGCTGGAAA
ATCTGATCGCCCAGCTGCCCGGCGAGAAGAAGAATGGCCTGTTCGGAAACCTGATTGCCCTGAGCC
TGGGCCTGACCCCCAACTTCAAGAGCAACTTCGACCTGGCCGAGGATGCCAAACTGCAGCTGAGCA
AGGACACCTACGACGACGACCTGGACAACCTGCTGGCCCAGATCGGCGACCAGTACGCCGACCTG
TTTCTGGCCGCCAAGAACCTGTCCGACGCCATCCTGCTGAGCGACATCCTGAGAGTGAACACCGAG
ATCACCAAGGCCCCCCTGAGCGCCTCTATGATCAAGAGATACGACGAGCACCACCAGGACCTGACC
CTGCTGAAAGCTCTCGTGCGGCAGCAGCTGCCTGAGAAGTACAAAGAGATTTTCTTCGACCAGAGC
AAGAACGGCTACGCCGGCTACATTGACGGCGGAGCCAGCCAGGAAGAGTTCTACAAGTTCATCAAG
CCCATCCTGGAAAAGATGGACGGCACCGAGGAACTGCTCGTGAAGCTGAACAGAGAGGACCTGCTG
CGGAAGCAGCGGACCTTCGACAACGGCAGCATCCCCCACCAGATCCACCTGGGAGAGCTGCACGC
CATTCTGCGGCGGCAGGAAGATTTTTACCCATTCCTGAAGGACAACCGGGAAAAGATCGAGAAGATC
CTGACCTTCCGCATCCCCTACTACGTGGGCCCTCTGGCCAGGGGAAACAGCAGATTCGCCTGGATG
ACCAGAAAGAGCGAGGAAACCATCACCCCCTGGAACTTCGAGGAAGTGGTGGACAAGGGCGCTTCC
GCCCAGAGCTTCATCGAGCGGATGACCAACTTCGATAAGAACCTGCCCAACGAGAAGGTGCTGCCC
AAGCACAGCCTGCTGTACGAGTACTTCACCGTGTATAACGAGCTGACCAAAGTGAAATACGTGACCG
AGGGAATGAGAAAGCCCGCCTTCCTGAGCGGCGAGCAGAAAAAGGCCATCGTGGACCTGCTGTTCA
AGACCAACCGGAAAGTGACCGTGAAGCAGCTGAAAGAGGACTACTTCAAGAAAATCGAGTGCTTCG
ACTCCGTGGAAATCTCCGGCGTGGAAGATCGGTTCAACGCCTCCCTGGGCACATACCACGATCTGC
TGAAAATTATCAAGGACAAGGACTTCCTGGACAATGAGGAAAACGAGGACATTCTGGAAGATATCGT
GCTGACCCTGACACTGTTTGAGGACAGAGAGATGATCGAGGAACGGCTGAAAACCTATGCCCACCT
GTTCGACGACAAAGTGATGAAGCAGCTGAAGCGGCGGAGATACACCGGCTGGGGCAGGCTGAGCC
GGAAGCTGATCAACGGCATCCGGGACAAGCAGTCCGGCAAGACAATCCTGGATTTCCTGAAGTCCG
ACGGCTTCGCCAACAGAAACTTCATGCAGCTGATCCACGACGACAGCCTGACCTTTAAAGAGGACAT
CCAGAAAGCCCAGGTGTCCGGCCAGGGCGATAGCCTGCACGAGCACATTGCCAATCTGGCCGGCA
GCCCCGCCATTAAGAAGGGCATCCTGCAGACAGTGAAGGTGGTGGACGAGCTCGTGAAAGTGATG
GGCCGGCACAAGCCCGAGAACATCGTGATCGAAATGGCCAGAGAGAACCAGACCACCCAGAAGGG
ACAGAAGAACAGCCGCGAGAGAATGAAGCGGATCGAAGAGGGCATCAAAGAGCTGGGCAGCCAGA
TCCTGAAAGAACACCCCGTGGAAAACACCCAGCTGCAGAACGAGAAGCTGTACCTGTACTACCTGC
AGAATGGGCGGGATATGTACGTGGACCAGGAACTGGACATCAACCGGCTGTCCGACTACGATGTGG
ACGCTATCGTGCCTCAGAGCTTTCTGGCCGACGACTCCATCGACAACAAGGTGCTGACCAGAAGCG

ACAAGAACCGGGGCAAGAGCGACAACGTGCCCTCCGAAGAGGTCGTGAAGAAGATGAAGAACTACT
GGCGGCAGCTGCTGAACGCCAAGCTGATTACCCAGAGAAAGTTCGACAATCTGACCAAGGCCGAGA
GAGGCGGCCTGAGCGAACTGGATAAGGCCGGCTTCATCAAGAGACAGCTGGTGGAAACCCGGCAG
ATCACAAAGCACGTGGCACAGATCCTGGACTCCCGGATGAACACTAAGTACGACGAGAATGACAAG
CTGATCCGGGAAGTGAAAGTGATCACCCTGAAGTCCAAGCTGGTGTCCGATTTCCGGAAGGATTTCC
AGTTTTACAAAGTGCGCGAGATCAACAACTACGCCCACGCCCACGACGCCTACCTGAACGCCGTCG
TGGGAACCGCCCTGATCAAAAAGTACCCTAAGCTGGAAAGCGAGTTCGTGTACGGCGACTACAAGG
TGTACGACGTGCGGAAGATGATCGCCAAGAGCGAGCAGGAAATCGGCAAGGCTACCGCCAAGTACT
TCTTCTACAGCAACATCATGAACTTTTTCAAGACCGAGATTACCCTGGCCAACGGCGAGATCCGGAA
GCGGCCTCTGATCGAGACAAACGGCGAAACCGGGGAGATCGTGTGGGATAAGGGCCGGGATTTTG
CCACCGTGCGGAAAGTGCTGAGCATGCCCCAAGTGAATATCGTGAAAAAGACCGAGGTGCAGACAG
GCGGCTTCAGCAAAGAGTCTATCCTGCCCAAGAGGAACAGCGATAAGCTGATCGCCAGAAAGAAGG
ACTGGGACCCTAAGAAGTACGGCGGCTTCGACAGCCCCACCGTGGCCTATTCTGTGCTGGTGGTGG
CCAAAGTGGAAAAGGGCAAGTCCAAGAAACTGAAGAGTGTGAAAGAGCTGCTGGGGATCACCATCA
TGGAAAGAAGCAGCTTCGAGAAGAATCCCATCGACTTTCTGGAAGCCAAGGGCTACAAAGAAGTGAA
AAAGGACCTGATCATCAAGCTGCCTAAGTACTCCCTGTTCGAGCTGGAAAACGGCCGGAAGAGAAT
GCTGGCCTCTGCCGGCGAACTGCAGAAGGGAAACGAACTGGCCCTGCCCTCCAAATATGTGAACTT
CCTGTACCTGGCCAGCCACTATGAGAAGCTGAAGGGCTCCCCCGAGGATAATGAGCAGAAACAGCT
GTTTGTGGAACAGCACAAGCACTACCTGGACGAGATCATCGAGCAGATCAGCGAGTTCTCCAAGAG
AGTGATCCTGGCCGACGCTAATCTGGACAAAGTGCTGTCCGCCTACAACAAGCACCGGGATAAGCC
CATCAGAGAGCAGGCCGAGAATATCATCCACCTGTTTACCCTGACCAATCTGGGAGCCCCTGCCGC
CTTCAAGTACTTTGACACCACCATCGACCGGAAGAGGTACACCAGCACCAAAGAGGTGCTGGACGC
CACCCTGATCCACCAGAGCATCACCGGCCTGTACGAGACACGGATCGACCTGTCTCAGCTGGGAGG
TGACTCCGGCGGAAGCTCTGGTGGCAGCAAGCGGACCGCCGACGGCTCTGAATTCGAGAGCCCTA
AGAAGAAAAGAAAGGTGAGCGGAGGCTCTAGCGGCGGAAGCACCCTGAACATTGAAGACGAGTATA
GACTGCATGAAACAAGCAAGGAACCCGACGTGTCCCTGGGCTCCACCTGGCTGTCCGACTTTCCCC
AGGCCTGGGCCGAGACAGGAGGAATGGGCCTGGCCGTGCGGCAGGCACCCCTGATCATCCCTCTG
AAGGCCACCTCTACACCCGTGAGCATCAAGCAGTACCCTATGTCTCAGGAGGCCAGACTGGGCATC
AAGCCTCACATCCAGAGGCTGCTGGACCAGGGCATCCTGGTGCCATGCCAGAGCCCCTGGAACACA
CCACTGCTGCCCGTGAAGAAGCCAGGCACCAATGACTATAGACCCGTGCAGGATCTGAGAGAGGTG
AACAAGAGGGTGGAGGATATCCACCCCACCGTGCCCAACCCTTACAATCTGCTGTCCGGCCTGCCC
CCTTCTCACCAGTGGTATACAGTGCTGGACCTGAAGGATGCCTTCTTTTGTCTGAGACTGCACCCTA
CCAGCCAGCCACTGTTCGCCTTTGAGTGGAGGGACCCTGAGATGGGCATCTCTGGCCAGCTGACCT
GGACACGCCTGCCTCAGGGCTTCAAGAATAGCCCAACACTGTTTAACGAGGCCCTGCACCGCGACC
TGGCAGATTTCCGGATCCAGCACCCAGATCTGATCCTGCTGCAGTACGTGGACGATCTGCTGCTGG
CCGCCACCAGCGAGCTGGATTGCCAGCAGGGAACACGCGCCCTGCTGCAGACCCTGGGAAACCTG

GGATATAGGGCATCCGCCAAGAAGGCCCAGATCTGTCAGAAGCAGGTGAAGTACCTGGGCTATCTG
CTGAAGGAGGGCCAGAGATGGCTGACAGAGGCCAGGAAGGAGACAGTGATGGGCCAGCCAACACC
CAAGACCCCAAGACAGCTGAGGGAGTTCCTGGGCAAAGCAGGATTTTGCAGGCTGTTCATCCCAGG
ATTCGCAGAGATGGCAGCACCTCTGTACCCACTGACCAAGCCGGGCACCCTGTTTAATTGGGGCCC
TGACCAGCAGAAGGCCTATCAGGAGATCAAGCAGGCCCTGCTGACAGCACCAGCCCTGGGCCTGC
CAGACCTGACCAAGCCTTTCGAGCTGTTTGTGGATGAGAAGCAGGGCTACGCCAAGGGCGTGCTGA
CCCAGAAGCTGGGACCATGGAGACGGCCCGTGGCCTATCTGTCCAAGAAGCTGGACCCAGTGGCA
GCAGGATGGCCACCATGCCTGAGGATGGTGGCAGCAATCGCCGTGCTGACAAAGGATGCCGGCAA
GCTGACCATGGGACAGCCACTGGTCATCCTGGCACCACACGCAGTGGAGGCCCTGGTGAAGCAGC
CTCCAGATCGCTGGCTGTCTAACGCCCGGATGACACACTACCAGGCCCTGCTGCTGGACACCGATC
GCGTGCAGTTTGGCCCTGTGGTGGCCCTGAATCCAGCCACCCTGCTGCCTCTGCCAGAGGAGGGC
CTGCAGCACAACTGTCTGGACATCCTGGCAGAGGCACACGGAACAAGGCCAGACCTGACCGATCAG
CCCCTGCCTGACGCCGATCACACATGGTATACCGATGGAAGCTCCCTGCTGCAGGAGGGCCAGAG
GAAGGCAGGAGCAGCAGTGACCACAGAGACAGAAGTGATCTGGGCCAAGGCCCTGCCAGCAGGCA
CATCCGCCCAGCGGGCCGAGCTGATCGCCCTGACCCAGGCCCTGAAGATGGCCGAGGGCAAGAAG
CTGAACGTGTACACAGACTCCAGATATGCCTTCGCCACCGCACACATCCACGGAGAGATCTACAGG
CGCCGGGGCTGGCTGACCTCTGAGGGCAAGGAGATCAAGAACAAGGATGAGATCCTGGCCCTGCT
GAAGGCCCTGTTTCTGCCCAAGCGGCTGAGCATCATCCACTGTCCTGGACACCAGAAGGGACACTC
CGCCGAGGCAAGGGGCAATCGGATGGCCGACCAGGCCGCCAGAAAGGCTGCTATTACTGAAACTC
CCGACACTTCCACTCTGCTGATTGAAAACTCCTCCCCTTCTGGCGGCTCAAAAAGAACCGCCGACGG
CAGCGAATTCGAGTCTCCCAAGAAGAAGAGGAAAGTCGGCTCTGGCCCTGCCGCTAAGAGAGTGAA
GCTGGAC

PE$^{R221K-K848A-H982A-N1317R}$ (xPE), with mutations (versus wild-type Cas9) in red:

ATGAAACGGACAGCCGACGGAAGCGAGTTCGAGTCACCAAAGAAGAAGCGGAAAGTCGACAAGAAG
TACAGCATCGGCCTGGACATCGGCACCAACTCTGTGGGCTGGGCCGTGATCACCGACGAGTACAAG
GTGCCCAGCAAGAAATTCAAGGTGCTGGGCAACACCGACCGGCACAGCATCAAGAAGAACCTGATC
GGAGCCCTGCTGTTCGACAGCGGCGAAACAGCCGAGGCCACCCGGCTGAAGAGAACCGCCAGAAG
AAGATACACCAGACGGAAGAACCGGATCTGCTATCTGCAAGAGATCTTCAGCAACGAGATGGCCAA
GGTGGACGACAGCTTCTTCCACAGACTGGAAGAGTCCTTCCTGGTGGAAGAGGATAAGAAGCACGA
GCGGCACCCCATCTTCGGCAACATCGTGGACGAGGTGGCCTACCACGAGAAGTACCCCACCATCTA
CCACCTGAGAAAGAAACTGGTGGACAGCACCGACAAGGCCGACCTGCGGCTGATCTATCTGGCCCT
GGCCCACATGATCAAGTTCCGGGGCCACTTCCTGATCGAGGGCGACCTGAACCCCGACAACAGCGA
CGTGGACAAGCTGTTCATCCAGCTGGTGCAGACCTACAACCAGCTGTTCGAGGAAAACCCCATCAA

CGCCAGCGGCGTGGACGCCAAGGCCATCCTGTCTGCCAGACTGAGCAAGAGCAGAAAGCTGGAAA
ATCTGATCGCCCAGCTGCCCGGCGAGAAGAAGAATGGCCTGTTCGGAAACCTGATTGCCCTGAGCC
TGGGCCTGACCCCCAACTTCAAGAGCAACTTCGACCTGGCCGAGGATGCCAAACTGCAGCTGAGCA
AGGACACCTACGACGACGACCTGGACAACCTGCTGGCCCAGATCGGCGACCAGTACGCCGACCTG
TTTCTGGCCGCCAAGAACCTGTCCGACGCCATCCTGCTGAGCGACATCCTGAGAGTGAACACCGAG
ATCACCAAGGCCCCCCTGAGCGCCTCTATGATCAAGAGATACGACGAGCACCACCAGGACCTGACC
CTGCTGAAAGCTCTCGTGCGGCAGCAGCTGCCTGAGAAGTACAAAGAGATTTTCTTCGACCAGAGC
AAGAACGGCTACGCCGGCTACATTGACGGCGGAGCCAGCCAGGAAGAGTTCTACAAGTTCATCAAG
CCCATCCTGGAAAAGATGGACGGCACCGAGGAACTGCTCGTGAAGCTGAACAGAGAGGACCTGCTG
CGGAAGCAGCGGACCTTCGACAACGGCAGCATCCCCCACCAGATCCACCTGGGAGAGCTGCACGC
CATTCTGCGGCGGCAGGAAGATTTTTACCCATTCCTGAAGGACAACCGGGAAAAGATCGAGAAGATC
CTGACCTTCCGCATCCCCTACTACGTGGGCCCTCTGGCCAGGGGAAACAGCAGATTCGCCTGGATG
ACCAGAAAGAGCGAGGAAACCATCACCCCCTGGAACTTCGAGGAAGTGGTGGACAAGGGCGCTTCC
GCCCAGAGCTTCATCGAGCGGATGACCAACTTCGATAAGAACCTGCCCAACGAGAAGGTGCTGCCC
AAGCACAGCCTGCTGTACGAGTACTTCACCGTGTATAACGAGCTGACCAAAGTGAAATACGTGACCG
AGGGAATGAGAAAGCCCGCCTTCCTGAGCGGCGAGCAGAAAAAGGCCATCGTGGACCTGCTGTTCA
AGACCAACCGGAAAGTGACCGTGAAGCAGCTGAAAGAGGACTACTTCAAGAAAATCGAGTGCTTCG
ACTCCGTGGAAATCTCCGGCGTGGAAGATCGGTTCAACGCCTCCCTGGGCACATACCACGATCTGC
TGAAAATTATCAAGGACAAGGACTTCCTGGACAATGAGGAAAACGAGGACATTCTGGAAGATATCGT
GCTGACCCTGACACTGTTTGAGGACAGAGAGATGATCGAGGAACGGCTGAAAACCTATGCCCACCT
GTTCGACGACAAAGTGATGAAGCAGCTGAAGCGGCGGAGATACACCGGCTGGGGCAGGCTGAGCC
GGAAGCTGATCAACGGCATCCGGGACAAGCAGTCCGGCAAGACAATCCTGGATTTCCTGAAGTCCG
ACGGCTTCGCCAACAGAAACTTCATGCAGCTGATCCACGACGACAGCCTGACCTTTAAAGAGGACAT
CCAGAAAGCCCAGGTGTCCGGCCAGGGCGATAGCCTGCACGAGCACATTGCCAATCTGGCCGGCA
GCCCCGCCATTAAGAAGGGCATCCTGCAGACAGTGAAGGTGGTGGACGAGCTCGTGAAAGTGATG
GGCCGGCACAAGCCCGAGAACATCGTGATCGAAATGGCCAGAGAGAACCAGACCACCCAGAAGGG
ACAGAAGAACAGCCGCGAGAGAATGAAGCGGATCGAAGAGGGCATCAAAGAGCTGGGCAGCCAGA
TCCTGAAAGAACACCCCGTGGAAAACACCCAGCTGCAGAACGAGAAGCTGTACCTGTACTACCTGC
AGAATGGGCGGGATATGTACGTGGACCAGGAACTGGACATCAACCGGCTGTCCGACTACGATGTGG
ACGCTATCGTGCCTCAGAGCTTTCTGGCCGACGACTCCATCGACAACAAGGTGCTGACCAGAAGCG
ACAAGAACCGGGGCAAGAGCGACAACGTGCCCTCCGAAGAGGTCGTGAAGAAGATGAAGAACTACT
GGCGGCAGCTGCTGAACGCCAAGCTGATTACCCAGAGAAAGTTCGACAATCTGACCAAGGCCGAGA
GAGGCGGCCTGAGCGAACTGGATAAGGCCGGCTTCATCAAGAGACAGCTGGTGGAAACCCGGCAG
ATCACAAAGCACGTGGCACAGATCCTGGACTCCCGGATGAACACTAAGTACGACGAGAATGACAAG
CTGATCCGGGAAGTGAAAGTGATCACCCTGAAGTCCAAGCTGGTGTCCGATTTCCGGAAGGATTTCC
AGTTTTACAAAGTGCGCGAGATCAACAACTACGCCCACGCCCACGACGCCTACCTGAACGCCGTCG

TGGGAACCGCCCTGATCAAAAAGTACCCTAAGCTGGAAAGCGAGTTCGTGTACGGCGACTACAAGG
TGTACGACGTGCGGAAGATGATCGCCAAGAGCGAGCAGGAAATCGGCAAGGCTACCGCCAAGTACT
TCTTCTACAGCAACATCATGAACTTTTTCAAGACCGAGATTACCCTGGCCAACGGCGAGATCCGGAA
GCGGCCTCTGATCGAGACAAACGGCGAAACCGGGGAGATCGTGTGGGATAAGGGCCGGGATTTTG
CCACCGTGCGGAAAGTGCTGAGCATGCCCCAAGTGAATATCGTGAAAAAGACCGAGGTGCAGACAG
GCGGCTTCAGCAAAGAGTCTATCCTGCCCAAGAGGAACAGCGATAAGCTGATCGCCAGAAAGAAGG
ACTGGGACCCTAAGAAGTACGGCGGCTTCGACAGCCCCACCGTGGCCTATTCTGTGCTGGTGGTGG
CCAAAGTGGAAAAGGGCAAGTCCAAGAAACTGAAGAGTGTGAAAGAGCTGCTGGGGATCACCATCA
TGGAAAGAAGCAGCTTCGAGAAGAATCCCATCGACTTTCTGGAAGCCAAGGGCTACAAAGAAGTGAA
AAAGGACCTGATCATCAAGCTGCCTAAGTACTCCCTGTTCGAGCTGGAAAACGGCCGGAAGAGAAT
GCTGGCCTCTGCCGGCGAACTGCAGAAGGGAAACGAACTGGCCCTGCCCTCCAAATATGTGAACTT
CCTGTACCTGGCCAGCCACTATGAGAAGCTGAAGGGCTCCCCCGAGGATAATGAGCAGAAACAGCT
GTTTGTGGAACAGCACAAGCACTACCTGGACGAGATCATCGAGCAGATCAGCGAGTTCTCCAAGAG
AGTGATCCTGGCCGACGCTAATCTGGACAAAGTGCTGTCCGCCTACAACAAGCACCGGGATAAGCC
CATCAGAGAGCAGGCCGAGAATATCATCCACCTGTTTACCCTGACCCGGCTGGGAGCCCCTGCCGC
CTTCAAGTACTTTGACACCACCATCGACCGGAAGAGGTACACCAGCACCAAAGAGGTGCTGGACGC
CACCCTGATCCACCAGAGCATCACCGGCCTGTACGAGACACGGATCGACCTGTCTCAGCTGGGAGG
TGACTCCGGCGGAAGCTCTGGTGGCAGCAAGCGGACCGCCGACGGCTCTGAATTCGAGAGCCCTA
AGAAGAAAAGAAAGGTGAGCGGAGGCTCTAGCGGCGGAAGCACCCTGAACATTGAAGACGAGTATA
GACTGCATGAAACAAGCAAGGAACCCGACGTGTCCCTGGGCTCCACCTGGCTGTCCGACTTTCCCC
AGGCCTGGGCCGAGACAGGAGGAATGGGCCTGGCCGTGCGGCAGGCACCCCTGATCATCCCTCTG
AAGGCCACCTCTACACCCGTGAGCATCAAGCAGTACCCTATGTCTCAGGAGGCCAGACTGGGCATC
AAGCCTCACATCCAGAGGCTGCTGGACCAGGGCATCCTGGTGCCATGCCAGAGCCCCTGGAACACA
CCACTGCTGCCCGTGAAGAAGCCAGGCACCAATGACTATAGACCCGTGCAGGATCTGAGAGAGGTG
AACAAGAGGGTGGAGGATATCCACCCCACCGTGCCCAACCCTTACAATCTGCTGTCCGGCCTGCCC
CCTTCTCACCAGTGGTATACAGTGCTGGACCTGAAGGATGCCTTCTTTTGTCTGAGACTGCACCCTA
CCAGCCAGCCACTGTTCGCCTTTGAGTGGAGGGACCCTGAGATGGGCATCTCTGGCCAGCTGACCT
GGACACGCCTGCCTCAGGGCTTCAAGAATAGCCCAACACTGTTTAACGAGGCCCTGCACCGCGACC
TGGCAGATTTCCGGATCCAGCACCCAGATCTGATCCTGCTGCAGTACGTGGACGATCTGCTGCTGG
CCGCCACCAGCGAGCTGGATTGCCAGCAGGGAACACGCGCCCTGCTGCAGACCCTGGGAAACCTG
GGATATAGGGCATCCGCCAAGAAGGCCCAGATCTGTCAGAAGCAGGTGAAGTACCTGGGCTATCTG
CTGAAGGAGGGCCAGAGATGGCTGACAGAGGCCAGGAAGGAGACAGTGATGGGCCAGCCAACACC
CAAGACCCCAAGACAGCTGAGGGAGTTCCTGGGCAAAGCAGGATTTTGCAGGCTGTTCATCCCAGG
ATTCGCAGAGATGGCAGCACCTCTGTACCCACTGACCAAGCCGGGCACCCTGTTTAATTGGGGCCC
TGACCAGCAGAAGGCCTATCAGGAGATCAAGCAGGCCCTGCTGACAGCACCAGCCCTGGGCCTGC
CAGACCTGACCAAGCCTTTCGAGCTGTTTGTGGATGAGAAGCAGGGCTACGCCAAGGGCGTGCTGA

CCCAGAAGCTGGGACCATGGAGACGGCCCGTGGCCTATCTGTCCAAGAAGCTGGACCCAGTGGCA
GCAGGATGGCCACCATGCCTGAGGATGGTGGCAGCAATCGCCGTGCTGACAAAGGATGCCGGCAA
GCTGACCATGGGACAGCCACTGGTCATCCTGGCACCACACGCAGTGGAGGCCCTGGTGAAGCAGC
CTCCAGATCGCTGGCTGTCTAACGCCCGGATGACACACTACCAGGCCCTGCTGCTGGACACCGATC
GCGTGCAGTTTGGCCCTGTGGTGGCCCTGAATCCAGCCACCCTGCTGCCTCTGCCAGAGGAGGGC
CTGCAGCACAACTGTCTGGACATCCTGGCAGAGGCACACGGAACAAGGCCAGACCTGACCGATCAG
CCCCTGCCTGACGCCGATCACACATGGTATACCGATGGAAGCTCCCTGCTGCAGGAGGGCCAGAG
GAAGGCAGGAGCAGCAGTGACCACAGAGACAGAAGTGATCTGGGCCAAGGCCCTGCCAGCAGGCA
CATCCGCCCAGCGGGCCGAGCTGATCGCCCTGACCCAGGCCCTGAAGATGGCCGAGGGCAAGAAG
CTGAACGTGTACACAGACTCCAGATATGCCTTCGCCACCGCACACATCCACGGAGAGATCTACAGG
CGCCGGGGCTGGCTGACCTCTGAGGGCAAGGAGATCAAGAACAAGGATGAGATCCTGGCCCTGCT
GAAGGCCCTGTTTCTGCCCAAGCGGCTGAGCATCATCCACTGTCCTGGACACCAGAAGGGACACTC
CGCCGAGGCAAGGGGCAATCGGATGGCCGACCAGGCCGCCAGAAAGGCTGCTATTACTGAAACTC
CCGACACTTCCACTCTGCTGATTGAAAACTCCTCCCCTTCTGGCGGCTCAAAAAGAACCGCCGACGG
CAGCGAATTCGAGTCTCCCAAGAAGAAGAGGAAAGTCGGCTCTGGCCCTGCCGCTAAGAGAGTGAA
GCTGGAC

PE7, with mutations (versus wild-type Cas9) in red

ATGAAACGGACAGCCGACGGAAGCGAGTTCGAGTCACCAAAGAAGAAGCGGAAAGTCGACAAGAAG
TACAGCATCGGCCTGGACATCGGCACCAACTCTGTGGGCTGGGCCGTGATCACCGACGAGTACAAG
GTGCCCAGCAAGAAATTCAAGGTGCTGGGCAACACCGACCGGCACAGCATCAAGAAGAACCTGATC
GGAGCCCTGCTGTTCGACAGCGGCGAAACAGCCGAGGCCACCCGGCTGAAGAGAACCGCCAGAAG
AAGATACACCAGACGGAAGAACCGGATCTGCTATCTGCAAGAGATCTTCAGCAACGAGATGGCCAA
GGTGGACGACAGCTTCTTCCACAGACTGGAAGAGTCCTTCCTGGTGGAAGAGGATAAGAAGCACGA
GCGGCACCCCATCTTCGGCAACATCGTGGACGAGGTGGCCTACCACGAGAAGTACCCCACCATCTA
CCACCTGAGAAAGAAACTGGTGGACAGCACCGACAAGGCCGACCTGCGGCTGATCTATCTGGCCCT
GGCCCACATGATCAAGTTCCGGGGCCACTTCCTGATCGAGGGCGACCTGAACCCCGACAACAGCGA
CGTGGACAAGCTGTTCATCCAGCTGGTGCAGACCTACAACCAGCTGTTCGAGGAAAACCCCATCAA
CGCCAGCGGCGTGGACGCCAAGGCCATCCTGTCTGCCAGACTGAGCAAGAGCAGAAAGCTGGAAA
ATCTGATCGCCCAGCTGCCCGGCGAGAAGAAGAATGGCCTGTTCGGAAACCTGATTGCCCTGAGCC
TGGGCCTGACCCCCAACTTCAAGAGCAACTTCGACCTGGCCGAGGATGCCAAACTGCAGCTGAGCA
AGGACACCTACGACGACGACCTGGACAACCTGCTGGCCCAGATCGGCGACCAGTACGCCGACCTG
TTTCTGGCCGCCAAGAACCTGTCCGACGCCATCCTGCTGAGCGACATCCTGAGAGTGAACACCGAG
ATCACCAAGGCCCCCCTGAGCGCCTCTATGATCAAGAGATACGACGAGCACCACCAGGACCTGACC

CTGCTGAAAGCTCTCGTGCGGCAGCAGCTGCCTGAGAAGTACAAAGAGATTTTCTTCGACCAGAGC
AAGAACGGCTACGCCGGCTACATTGACGGCGGAGCCAGCCAGGAAGAGTTCTACAAGTTCATCAAG
CCCATCCTGGAAAAGATGGACGGCACCGAGGAACTGCTCGTGAAGCTGAAGAGAGAGGACCTGCT
GCGGAAGCAGCGGACCTTCGACAACGGCAGCATCCCCCACCAGATCCACCTGGGAGAGCTGCACG
CCATTCTGCGGCGGCAGGAAGATTTTTACCCATTCCTGAAGGACAACCGGGAAAAGATCGAGAAGAT
CCTGACCTTCCGCATCCCCTACTACGTGGGCCCTCTGGCCAGGGGAAACAGCAGATTCGCCTGGAT
GACCAGAAAGAGCGAGGAAACCATCACCCCCTGGAACTTCGAGGAAGTGGTGGACAAGGGCGCTT
CCGCCCAGAGCTTCATCGAGCGGATGACCAACTTCGATAAGAACCTGCCCAACGAGAAGGTGCTGC
CCAAGCACAGCCTGCTGTACGAGTACTTCACCGTGTATAACGAGCTGACCAAAGTGAAATACGTGAC
CGAGGGAATGAGAAAGCCCGCCTTCCTGAGCGGCGAGCAGAAAAAGGCCATCGTGGACCTGCTGT
TCAAGACCAACCGGAAAGTGACCGTGAAGCAGCTGAAAGAGGACTACTTCAAGAAAATCGAGTGCTT
CGACTCCGTGGAAATCTCCGGCGTGGAAGATCGGTTCAACGCCTCCCTGGGCACATACCACGATCT
GCTGAAAATTATCAAGGACAAGGACTTCCTGGACAATGAGGAAAACGAGGACATTCTGGAAGATATC
GTGCTGACCCTGACACTGTTTGAGGACAGAGAGATGATCGAGGAACGGCTGAAAACCTATGCCCAC
CTGTTCGACGACAAAGTGATGAAGCAGCTGAAGCGGCGGAGATACACCGGCTGGGGCAGGCTGAG
CCGGAAGCTGATCAACGGCATCCGGGACAAGCAGTCCGGCAAGACAATCCTGGATTTCCTGAAGTC
CGACGGCTTCGCCAACAGAAACTTCATGCAGCTGATCCACGACGACAGCCTGACCTTTAAAGAGGA
CATCCAGAAAGCCCAGGTGTCCGGCCAGGGCGATAGCCTGCACGAGCACATTGCCAATCTGGCCG
GCAGCCCCGCCATTAAGAAGGGCATCCTGCAGACAGTGAAGGTGGTGGACGAGCTCGTGAAAGTG
ATGGGCCGGCACAAGCCCGAGAACATCGTGATCGAAATGGCCAGAGAGAACCAGACCACCCAGAA
GGGACAGAAGAACAGCCGCGAGAGAATGAAGCGGATCGAAGAGGGCATCAAAGAGCTGGGCAGCC
AGATCCTGAAAGAACACCCCGTGGAAAACACCCAGCTGCAGAACGAGAAGCTGTACCTGTACTACCT
GCAGAATGGGCGGGATATGTACGTGGACCAGGAACTGGACATCAACCGGCTGTCCGACTACGATGT
GGACGCTATCGTGCCTCAGAGCTTTCTGAAGGACGACTCCATCGACAACAAGGTGCTGACCAGAAG
CGACAAGAACCGGGGCAAGAGCGACAACGTGCCCTCCGAAGAGGTCGTGAAGAAGATGAAGAACT
ACTGGCGGCAGCTGCTGAACGCCAAGCTGATTACCCAGAGAAAGTTCGACAATCTGACCAAGGCCG
AGAGAGGCGGCCTGAGCGAACTGGATAAGGCCGGCTTCATCAAGAGACAGCTGGTGGAAACCCGG
CAGATCACAAAGCACGTGGCACAGATCCTGGACTCCCGGATGAACACTAAGTACGACGAGAATGAC
AAGCTGATCCGGGAAGTGAAAGTGATCACCCTGAAGTCCAAGCTGGTGTCCGATTTCCGGAAGGAT
TTCCAGTTTTACAAAGTGCGCGAGATCAACAACTACCACCACGCCCACGACGCCTACCTGAACGCCG
TCGTGGGAACCGCCCTGATCAAAAAGTACCCTAAGCTGGAAAGCGAGTTCGTGTACGGCGACTACA
AGGTGTACGACGTGCGGAAGATGATCGCCAAGAGCGAGCAGGAAATCGGCAAGGCTACCGCCAAG
TACTTCTTCTACAGCAACATCATGAACTTTTTCAAGACCGAGATTACCCTGGCCAACGGCGAGATCC
GGAAGCGGCCTCTGATCGAGACAAACGGCGAAACCGGGGAGATCGTGTGGGATAAGGGCCGGGAT
TTTGCCACCGTGCGGAAAGTGCTGAGCATGCCCCAAGTGAATATCGTGAAAAAGACCGAGGTGCAG
ACAGGCGGCTTCAGCAAAGAGTCTATCCTGCCCAAGAGGAACAGCGATAAGCTGATCGCCAGAAAG

AAGGACTGGGACCCTAAGAAGTACGGCGGCTTCGACAGCCCCACCGTGGCCTATTCTGTGCTGGTG
GTGGCCAAAGTGGAAAAGGGCAAGTCCAAGAAACTGAAGAGTGTGAAAGAGCTGCTGGGGATCACC
ATCATGGAAAGAAGCAGCTTCGAGAAGAATCCCATCGACTTTCTGGAAGCCAAGGGCTACAAAGAAG
TGAAAAAGGACCTGATCATCAAGCTGCCTAAGTACTCCCTGTTCGAGCTGGAAAACGGCCGGAAGA
GAATGCTGGCCTCTGCCGGCGAACTGCAGAAGGGAAACGAACTGGCCCTGCCCTCCAAATATGTGA
ACTTCCTGTACCTGGCCAGCCACTATGAGAAGCTGAAGGGCTCCCCCGAGGATAATGAGCAGAAAC
AGCTGTTTGTGGAACAGCACAAGCACTACCTGGACGAGATCATCGAGCAGATCAGCGAGTTCTCCAA
GAGAGTGATCCTGGCCGACGCTAATCTGGACAAAGTGCTGTCCGCCTACAACAAGCACCGGGATAA
GCCCATCAGAGAGCAGGCCGAGAATATCATCCACCTGTTTACCCTGACCAATCTGGGAGCCCCTGC
CGCCTTCAAGTACTTTGACACCACCATCGACCGGAAGAGGTACACCAGCACCAAAGAGGTGCTGGA
CGCCACCCTGATCCACCAGAGCATCACCGGCCTGTACGAGACACGGATCGACCTGTCTCAGCTGGG
AGGTGACTCCGGCGGAAGCTCTGGTGGCAGCAAGCGGACCGCCGACGGCTCTGAATTCGAGAGCC
CTAAGAAGAAAAGAAAGGTGAGCGGAGGCTCTAGCGGCGGAAGCACCCTGAACATTGAAGACGAGT
ATAGACTGCATGAAACAAGCAAGGAACCCGACGTGTCCCTGGGCTCCACCTGGCTGTCCGACTTTC
CCCAGGCCTGGGCCGAGACAGGAGGAATGGGCCTGGCCGTGCGGCAGGCACCCCTGATCATCCCT
CTGAAGGCCACCTCTACACCCGTGAGCATCAAGCAGTACCCTATGTCTCAGGAGGCCAGACTGGGC
ATCAAGCCTCACATCCAGAGGCTGCTGGACCAGGGCATCCTGGTGCCATGCCAGAGCCCCTGGAAC
ACACCACTGCTGCCCGTGAAGAAGCCAGGCACCAATGACTATAGACCCGTGCAGGATCTGAGAGAG
GTGAACAAGAGGGTGGAGGATATCCACCCCACCGTGCCCAACCCTTACAATCTGCTGTCCGGCCTG
CCCCCTTCTCACCAGTGGTATACAGTGCTGGACCTGAAGGATGCCTTCTTTTGTCTGAGACTGCACC
CTACCAGCCAGCCACTGTTCGCCTTTGAGTGGAGGGACCCTGAGATGGGCATCTCTGGCCAGCTGA
CCTGGACACGCCTGCCTCAGGGCTTCAAGAATAGCCCAACACTGTTTAACGAGGCCCTGCACCGCG
ACCTGGCAGATTTCCGGATCCAGCACCCAGATCTGATCCTGCTGCAGTACGTGGACGATCTGCTGC
TGGCCGCCACCAGCGAGCTGGATTGCCAGCAGGGAACACGCGCCCTGCTGCAGACCCTGGGAAAC
CTGGGATATAGGGCATCCGCCAAGAAGGCCCAGATCTGTCAGAAGCAGGTGAAGTACCTGGGCTAT
CTGCTGAAGGAGGGCCAGAGATGGCTGACAGAGGCCAGGAAGGAGACAGTGATGGGCCAGCCAAC
ACCCAAGACCCCAAGACAGCTGAGGGAGTTCCTGGGCAAAGCAGGATTTTGCAGGCTGTTCATCCC
AGGATTCGCAGAGATGGCAGCACCTCTGTACCCACTGACCAAGCCGGGCACCCTGTTTAATTGGGG
CCCTGACCAGCAGAAGGCCTATCAGGAGATCAAGCAGGCCCTGCTGACAGCACCAGCCCTGGGCC
TGCCAGACCTGACCAAGCCTTTCGAGCTGTTTGTGGATGAGAAGCAGGGCTACGCCAAGGGCGTGC
TGACCCAGAAGCTGGGACCATGGAGACGGCCCGTGGCCTATCTGTCCAAGAAGCTGGACCCAGTG
GCAGCAGGATGGCCACCATGCCTGAGGATGGTGGCAGCAATCGCCGTGCTGACAAAGGATGCCGG
CAAGCTGACCATGGGACAGCCACTGGTCATCCTGGCACCACACGCAGTGGAGGCCCTGGTGAAGC
AGCCTCCAGATCGCTGGCTGTCTAACGCCCGGATGACACACTACCAGGCCCTGCTGCTGGACACCG
ATCGCGTGCAGTTTGGCCCTGTGGTGGCCCTGAATCCAGCCACCCTGCTGCCTCTGCCAGAGGAGG
GCCTGCAGCACAACTGTCTGGACATCCTGGCAGAGGCACACGGAACAAGGCCAGACCTGACCGATC

AGCCCCTGCCTGACGCCGATCACACATGGTATACCGATGGAAGCTCCCTGCTGCAGGAGGGCCAGA
GGAAGGCAGGAGCAGCAGTGACCACAGAGACAGAAGTGATCTGGGCCAAGGCCCTGCCAGCAGGC
ACATCCGCCCAGCGGGCCGAGCTGATCGCCCTGACCCAGGCCCTGAAGATGGCCGAGGGCAAGAA
GCTGAACGTGTACACAGACTCCAGATATGCCTTCGCCACCGCACACATCCACGGAGAGATCTACAG
GCGCCGGGGCTGGCTGACCTCTGAGGGCAAGGAGATCAAGAACAAGGATGAGATCCTGGCCCTGC
TGAAGGCCCTGTTTCTGCCCAAGCGGCTGAGCATCATCCACTGTCCTGGACACCAGAAGGGACACT
CCGCCGAGGCAAGGGGCAATCGGATGGCCGACCAGGCCGCCAGAAAGGCTGCTATTACTGAAACT
CCCGACACTTCCACTCTGCTGATTGAAAACTCCTCCCCTTCCGGAGGATCTAGCGGAGGCTCCTCTG
GCTCTGAGACACCTGGCACAAGCGAGAGCGCAACACCTGAAAGCAGCGGGGGCAGCAGCGGGGG
GTCAATGGCTGAAAATGGTGATAATGAAAAGATGGCTGCCCTGGAGGCCAAAATCTGTCATCAAATT
GAGTATTATTTTGGCGACTTCAATTTGCCACGGGACAAGTTTCTAAAGGAACAGATAAAACTGGATGA
AGGCTGGGTACCTTTGGAGATAATGATAAAATTCAACAGGTTGAACCGTCTAACAACAGACTTTAATG
TAATTGTGGAAGCATTGAGCAAATCCAAGGCAGAACTCATGGAAATCAGTGAAGATAAAACTAAAATC
AGAAGGTCTCCAAGCAAACCCCTACCTGAAGTGACTGATGAGTATAAAAATGATGTAAAAAACAGATC
TGTTTATATTAAAGGCTTCCCAACTGATGCAACTCTTGATGACATAAAAGAATGGTTAGAAGATAAAG
GTCAAGTACTAAATATTCAGATGAGAAGAACATTGCATAAAGCATTTAAGGGATCAATTTTTGTTGTGT
TTGATAGCATTGAATCTGCTAAGAAATTTGTAGAGACCCCTGGCCAGAAGTACAAAGAAACAGACCT
GCTAATACTTTTCAAGGACGATTACTTTGCCAAAAAAAATGAATCTGGCGGCTCAAAAAGAACCGCCG
ACGGCAGCGAATTCGAGTCTCCCAAGAAGAAGAGGAAAGTCGGCTCTGGCCCTGCCGCTAAGAGAG
TGAAGCTGGAC

vPE, with mutations (versus wild-type Cas9) in red

ATGAAACGGACAGCCGACGGAAGCGAGTTCGAGTCACCAAAGAAGAAGCGGAAAGTCGACAAGAAG
TACAGCATCGGCCTGGACATCGGCACCAACTCTGTGGGCTGGGCCGTGATCACCGACGAGTACAAG
GTGCCCAGCAAGAAATTCAAGGTGCTGGGCAACACCGACCGGCACAGCATCAAGAAGAACCTGATC
GGAGCCCTGCTGTTCGACAGCGGCGAAACAGCCGAGGCCACCCGGCTGAAGAGAACCGCCAGAAG
AAGATACACCAGACGGAAGAACCGGATCTGCTATCTGCAAGAGATCTTCAGCAACGAGATGGCCAA
GGTGGACGACAGCTTCTTCCACAGACTGGAAGAGTCCTTCCTGGTGGAAGAGGATAAGAAGCACGA
GCGGCACCCCATCTTCGGCAACATCGTGGACGAGGTGGCCTACCACGAGAAGTACCCCACCATCTA
CCACCTGAGAAAGAAACTGGTGGACAGCACCGACAAGGCCGACCTGCGGCTGATCTATCTGGCCCT
GGCCCACATGATCAAGTTCCGGGGCCACTTCCTGATCGAGGGCGACCTGAACCCCGACAACAGCGA
CGTGGACAAGCTGTTCATCCAGCTGGTGCAGACCTACAACCAGCTGTTCGAGGAAAACCCCATCAA
CGCCAGCGGCGTGGACGCCAAGGCCATCCTGTCTGCCAGACTGAGCAAGAGCAGAAAGCTGGAAA
ATCTGATCGCCCAGCTGCCCGGCGAGAAGAAGAATGGCCTGTTCGGAAACCTGATTGCCCTGAGCC

TGGGCCTGACCCCCAACTTCAAGAGCAACTTCGACCTGGCCGAGGATGCCAAACTGCAGCTGAGCA
AGGACACCTACGACGACGACCTGGACAACCTGCTGGCCCAGATCGGCGACCAGTACGCCGACCTG
TTTCTGGCCGCCAAGAACCTGTCCGACGCCATCCTGCTGAGCGACATCCTGAGAGTGAACACCGAG
ATCACCAAGGCCCCCCTGAGCGCCTCTATGATCAAGAGATACGACGAGCACCACCAGGACCTGACC
CTGCTGAAAGCTCTCGTGCGGCAGCAGCTGCCTGAGAAGTACAAAGAGATTTTCTTCGACCAGAGC
AAGAACGGCTACGCCGGCTACATTGACGGCGGAGCCAGCCAGGAAGAGTTCTACAAGTTCATCAAG
CCCATCCTGGAAAAGATGGACGGCACCGAGGAACTGCTCGTGAAGCTGAACAGAGAGGACCTGCTG
CGGAAGCAGCGGACCTTCGACAACGGCAGCATCCCCCACCAGATCCACCTGGGAGAGCTGCACGC
CATTCTGCGGCGGCAGGAAGATTTTTACCCATTCCTGAAGGACAACCGGGAAAAGATCGAGAAGATC
CTGACCTTCCGCATCCCCTACTACGTGGGCCCTCTGGCCAGGGGAAACAGCAGATTCGCCTGGATG
ACCAGAAAGAGCGAGGAAACCATCACCCCCTGGAACTTCGAGGAAGTGGTGGACAAGGGCGCTTCC
GCCCAGAGCTTCATCGAGCGGATGACCAACTTCGATAAGAACCTGCCCAACGAGAAGGTGCTGCCC
AAGCACAGCCTGCTGTACGAGTACTTCACCGTGTATAACGAGCTGACCAAAGTGAAATACGTGACCG
AGGGAATGAGAAAGCCCGCCTTCCTGAGCGGCGAGCAGAAAAAGGCCATCGTGGACCTGCTGTTCA
AGACCAACCGGAAAGTGACCGTGAAGCAGCTGAAAGAGGACTACTTCAAGAAAATCGAGTGCTTCG
ACTCCGTGGAAATCTCCGGCGTGGAAGATCGGTTCAACGCCTCCCTGGGCACATACCACGATCTGC
TGAAAATTATCAAGGACAAGGACTTCCTGGACAATGAGGAAAACGAGGACATTCTGGAAGATATCGT
GCTGACCCTGACACTGTTTGAGGACAGAGAGATGATCGAGGAACGGCTGAAAACCTATGCCCACCT
GTTCGACGACAAAGTGATGAAGCAGCTGAAGCGGCGGAGATACACCGGCTGGGGCAGGCTGAGCC
GGAAGCTGATCAACGGCATCCGGGACAAGCAGTCCGGCAAGACAATCCTGGATTTCCTGAAGTCCG
ACGGCTTCGCCAACAGAAACTTCATGCAGCTGATCCACGACGACAGCCTGACCTTTAAAGAGGACAT
CCAGAAAGCCCAGGTGTCCGGCCAGGGCGATAGCCTGCACGAGCACATTGCCAATCTGGCCGGCA
GCCCCGCCATTAAGAAGGGCATCCTGCAGACAGTGAAGGTGGTGGACGAGCTCGTGAAAGTGATG
GGCCGGCACAAGCCCGAGAACATCGTGATCGAAATGGCCAGAGAGAACCAGACCACCCAGAAGGG
ACAGAAGAACAGCCGCGAGAGAATGAAGCGGATCGAAGAGGGCATCAAAGAGCTGGGCAGCCAGA
TCCTGAAAGAACACCCCGTGGAAAACACCCAGCTGCAGAACGAGAAGCTGTACCTGTACTACCTGC
AGAATGGGCGGGATATGTACGTGGACCAGGAACTGGACATCAACCGGCTGTCCGACTACGATGTGG
ACGCTATCGTGCCTCAGAGCTTTCTGGCCGACGACTCCATCGACAACAAGGTGCTGACCAGAAGCG
ACAAGAACCGGGGCAAGAGCGACAACGTGCCCTCCGAAGAGGTCGTGAAGAAGATGAAGAACTACT
GGCGGCAGCTGCTGAACGCCAAGCTGATTACCCAGAGAAAGTTCGACAATCTGACCAAGGCCGAGA
GAGGCGGCCTGAGCGAACTGGATAAGGCCGGCTTCATCAAGAGACAGCTGGTGGAAACCCGGCAG
ATCACAAAGCACGTGGCACAGATCCTGGACTCCCGGATGAACACTAAGTACGACGAGAATGACAAG
CTGATCCGGGAAGTGAAAGTGATCACCCTGAAGTCCAAGCTGGTGTCCGATTTCCGGAAGGATTTCC
AGTTTTACAAAGTGCGCGAGATCAACAACTACGCCCACGCCCACGACGCCTACCTGAACGCCGTCG
TGGGAACCGCCCTGATCAAAAAGTACCCTAAGCTGGAAAGCGAGTTCGTGTACGGCGACTACAAGG
TGTACGACGTGCGGAAGATGATCGCCAAGAGCGAGCAGGAAATCGGCAAGGCTACCGCCAAGTACT

TCTTCTACAGCAACATCATGAACTTTTTCAAGACCGAGATTACCCTGGCCAACGGCGAGATCCGGAA
GCGGCCTCTGATCGAGACAAACGGCGAAACCGGGGAGATCGTGTGGGATAAGGGCCGGGATTTTG
CCACCGTGCGGAAAGTGCTGAGCATGCCCCAAGTGAATATCGTGAAAAAGACCGAGGTGCAGACAG
GCGGCTTCAGCAAAGAGTCTATCCTGCCCAAGAGGAACAGCGATAAGCTGATCGCCAGAAAGAAGG
ACTGGGACCCTAAGAAGTACGGCGGCTTCGACAGCCCCACCGTGGCCTATTCTGTGCTGGTGGTGG
CCAAAGTGGAAAAGGGCAAGTCCAAGAAACTGAAGAGTGTGAAAGAGCTGCTGGGGATCACCATCA
TGGAAAGAAGCAGCTTCGAGAAGAATCCCATCGACTTTCTGGAAGCCAAGGGCTACAAAGAAGTGAA
AAAGGACCTGATCATCAAGCTGCCTAAGTACTCCCTGTTCGAGCTGGAAAACGGCCGGAAGAGAAT
GCTGGCCTCTGCCGGCGAACTGCAGAAGGGAAACGAACTGGCCCTGCCCTCCAAATATGTGAACTT
CCTGTACCTGGCCAGCCACTATGAGAAGCTGAAGGGCTCCCCCGAGGATAATGAGCAGAAACAGCT
GTTTGTGGAACAGCACAAGCACTACCTGGACGAGATCATCGAGCAGATCAGCGAGTTCTCCAAGAG
AGTGATCCTGGCCGACGCTAATCTGGACAAAGTGCTGTCCGCCTACAACAAGCACCGGGATAAGCC
CATCAGAGAGCAGGCCGAGAATATCATCCACCTGTTTACCCTGACCCGGCTGGGAGCCCCTGCCGC
CTTCAAGTACTTTGACACCACCATCGACCGGAAGAGGTACACCAGCACCAAAGAGGTGCTGGACGC
CACCCTGATCCACCAGAGCATCACCGGCCTGTACGAGACACGGATCGACCTGTCTCAGCTGGGAGG
TGACTCCGGCGGAAGCTCTGGTGGCAGCAAGCGGACCGCCGACGGCTCTGAATTCGAGAGCCCTA
AGAAGAAAAGAAAGGTGAGCGGAGGCTCTAGCGGCGGAAGCACCCTGAACATTGAAGACGAGTATA
GACTGCATGAAACAAGCAAGGAACCCGACGTGTCCCTGGGCTCCACCTGGCTGTCCGACTTTCCCC
AGGCCTGGGCCGAGACAGGAGGAATGGGCCTGGCCGTGCGGCAGGCACCCCTGATCATCCCTCTG
AAGGCCACCTCTACACCCGTGAGCATCAAGCAGTACCCTATGTCTCAGGAGGCCAGACTGGGCATC
AAGCCTCACATCCAGAGGCTGCTGGACCAGGGCATCCTGGTGCCATGCCAGAGCCCCTGGAACACA
CCACTGCTGCCCGTGAAGAAGCCAGGCACCAATGACTATAGACCCGTGCAGGATCTGAGAGAGGTG
AACAAGAGGGTGGAGGATATCCACCCCACCGTGCCCAACCCTTACAATCTGCTGTCCGGCCTGCCC
CCTTCTCACCAGTGGTATACAGTGCTGGACCTGAAGGATGCCTTCTTTTGTCTGAGACTGCACCCTA
CCAGCCAGCCACTGTTCGCCTTTGAGTGGAGGGACCCTGAGATGGGCATCTCTGGCCAGCTGACCT
GGACACGCCTGCCTCAGGGCTTCAAGAATAGCCCAACACTGTTTAACGAGGCCCTGCACCGCGACC
TGGCAGATTTCCGGATCCAGCACCCAGATCTGATCCTGCTGCAGTACGTGGACGATCTGCTGCTGG
CCGCCACCAGCGAGCTGGATTGCCAGCAGGGAACACGCGCCCTGCTGCAGACCCTGGGAAACCTG
GGATATAGGGCATCCGCCAAGAAGGCCCAGATCTGTCAGAAGCAGGTGAAGTACCTGGGCTATCTG
CTGAAGGAGGGCCAGAGATGGCTGACAGAGGCCAGGAAGGAGACAGTGATGGGCCAGCCAACACC
CAAGACCCCAAGACAGCTGAGGGAGTTCCTGGGCAAAGCAGGATTTTGCAGGCTGTTCATCCCAGG
ATTCGCAGAGATGGCAGCACCTCTGTACCCACTGACCAAGCCGGGCACCCTGTTTAATTGGGGCCC
TGACCAGCAGAAGGCCTATCAGGAGATCAAGCAGGCCCTGCTGACAGCACCAGCCCTGGGCCTGC
CAGACCTGACCAAGCCTTTCGAGCTGTTTGTGGATGAGAAGCAGGGCTACGCCAAGGGCGTGCTGA
CCCAGAAGCTGGGACCATGGAGACGGCCCGTGGCCTATCTGTCCAAGAAGCTGGACCCAGTGGCA
GCAGGATGGCCACCATGCCTGAGGATGGTGGCAGCAATCGCCGTGCTGACAAAGGATGCCGGCAA

GCTGACCATGGGACAGCCACTGGTCATCCTGGCACCACACGCAGTGGAGGCCCTGGTGAAGCAGC
CTCCAGATCGCTGGCTGTCTAACGCCCGGATGACACACTACCAGGCCCTGCTGCTGGACACCGATC
GCGTGCAGTTTGGCCCTGTGGTGGCCCTGAATCCAGCCACCCTGCTGCCTCTGCCAGAGGAGGGC
CTGCAGCACAACTGTCTGGACATCCTGGCAGAGGCACACGGAACAAGGCCAGACCTGACCGATCAG
CCCCTGCCTGACGCCGATCACACATGGTATACCGATGGAAGCTCCCTGCTGCAGGAGGGCCAGAG
GAAGGCAGGAGCAGCAGTGACCACAGAGACAGAAGTGATCTGGGCCAAGGCCCTGCCAGCAGGCA
CATCCGCCCAGCGGGCCGAGCTGATCGCCCTGACCCAGGCCCTGAAGATGGCCGAGGGCAAGAAG
CTGAACGTGTACACAGACTCCAGATATGCCTTCGCCACCGCACACATCCACGGAGAGATCTACAGG
CGCCGGGGCTGGCTGACCTCTGAGGGCAAGGAGATCAAGAACAAGGATGAGATCCTGGCCCTGCT
GAAGGCCCTGTTTCTGCCCAAGCGGCTGAGCATCATCCACTGTCCTGGACACCAGAAGGGACACTC
CGCCGAGGCAAGGGGCAATCGGATGGCCGACCAGGCCGCCAGAAAGGCTGCTATTACTGAAACTC
CCGACACTTCCACTCTGCTGATTGAAAACTCCTCCCCTTCCGGAGGATCTAGCGGAGGCTCCTCTGG
CTCTGAGACACCTGGCACAAGCGAGAGCGCAACACCTGAAAGCAGCGGGGGCAGCAGCGGGGGG
TCAATGGCTGAAAATGGTGATAATGAAAAGATGGCTGCCCTGGAGGCCAAAATCTGTCATCAAATTG
AGTATTATTTTGGCGACTTCAATTTGCCACGGGACAAGTTTCTAAAGGAACAGATAAAACTGGATGAA
GGCTGGGTACCTTTGGAGATAATGATAAAATTCAACAGGTTGAACCGTCTAACAACAGACTTTAATGT
AATTGTGGAAGCATTGAGCAAATCCAAGGCAGAACTCATGGAAATCAGTGAAGATAAAACTAAAATCA
GAAGGTCTCCAAGCAAACCCCTACCTGAAGTGACTGATGAGTATAAAAATGATGTAAAAAACAGATCT
GTTTATATTAAAGGCTTCCCAACTGATGCAACTCTTGATGACATAAAAGAATGGTTAGAAGATAAAGG
TCAAGTACTAAATATTCAGATGAGAAGAACATTGCATAAAGCATTTAAGGGATCAATTTTTGTTGTGTT
TGATAGCATTGAATCTGCTAAGAAATTTGTAGAGACCCCTGGCCAGAAGTACAAAGAAACAGACCTG
CTAATACTTTTCAAGGACGATTACTTTGCCAAAAAAAATGAATCTGGCGGCTCAAAAAGAACCGCCGA
CGGCAGCGAATTCGAGTCTCCCAAGAAGAAGAGGAAAGTCGGCTCTGGCCCTGCCGCTAAGAGAGT
GAAGCTGGAC
